# Integrative proteomics reveals principles of dynamic phospho-signaling networks in human erythropoiesis

**DOI:** 10.1101/2020.05.18.102178

**Authors:** Özge Karayel, Peng Xu, Isabell Bludau, Senthil Velan Bhoopalan, Yu Yao, Ana Rita Freitas Colaco, Alberto Santos Delgado, Brenda A. Schulman, Arno F. Alpi, Mitchell J. Weiss, Matthias Mann

## Abstract

Human erythropoiesis is exquisitely controlled at multiple levels and its dysregulation leads to numerous human diseases. Despite many functional studies focused on classical regulators, we lack a global, system-wide understanding of post-translational mechanisms coordinating erythroid maturation. Using the latest advances in mass spectrometry (MS)-based proteomics we comprehensively investigate the dynamics of protein and post-translational regulation of *in vitro* reconstituted CD34^+^ HSPC-derived erythropoiesis. This quantifies and dynamically tracks 7,400 proteins and 27,000 phosphorylation sites. Our data reveals differential temporal protein expression encompassing most protein classes and numerous post-translational regulatory cascades. Drastic cell surface remodeling across erythropoiesis include numerous orchestrated changes in solute carriers, providing new stage-specific markers. The dynamic phosphoproteomes combined with a kinome-targeting CRISPR/Cas9 screen reveal coordinated networks of erythropoietic kinases and downregulation of MAPK signaling subsequent to c-Kit attenuation as key drivers of maturation. Our global view of erythropoiesis establishes a central role of post-translational regulation in terminal differentiation.

## INTRODUCTION

Human erythropoiesis is a multistep developmental process that maintains stable erythroid homeostasis throughout life and replenishes more than 200 billion erythrocytes lost by senescence in healthy humans (Palis, 2014). Lineage-committed erythroid progenitors, including burst-forming unit-erythroid (BFU-E) and their colony-forming unit-erythroid, (CFU-E) progeny, undergo enormous expansion, followed by morphological signs of terminal maturation. The first recognizable erythroid precursors are proerythroblasts (ProE), which mature progressively into early basophilic (EBaso) and late basophilic (LBaso) erythroblasts, polychromatic erythroblasts (Poly), orthochromatic (Ortho) erythroblasts and reticulocytes. Terminal erythroid maturation is distinguished by progressive reductions in proliferative capacity and cell size, chromatin condensation, loss of most organelles including the nucleus, and remarkable streamlining of the proteome with expression of specialized cytoskeletal and plasma membrane proteins and finally massive accumulation of hemoglobin (Moras et al., 2017; Nguyen et al., 2017; Zhao et al., 2016b). This finely tuned developmental process generates mature erythrocytes with the highly specialized function of circulatory oxygen/carbon dioxide transport.

Our knowledge of human erythropoiesis has been greatly advanced by *in vitro* differentiation systems in which primary multipotent CD34^+^ hematopoietic stem cell progenitors (HSPCs) are cultured with defined cytokines and other bioactive components to generate reticulocytes (Seo et al., 2019). Erythropoiesis is controlled by the essential cytokines stem cell factor (SCF) and erythropoietin (EPO), and their cognate receptors c-Kit and EPOR (Broudy, 1997; Ingley, 2012; Nocka et al., 1989; Wu et al., 1997; Wu et al., 1995; Zhang and Lodish, 2008). In general, c-Kit acts to promote progenitor proliferation during early erythropoiesis, while EPOR fosters survival and maturation at later stages, although there is substantial overlap in their activities and some evidence for cross-regulation (Klingmuller, 1997; Wojchowski et al., 1999; Wu et al., 1997). Moreover, c-Kit and EPOR trigger remarkably similar signaling pathways including Ras/Raf/MAPK, PI3K/Akt, and JAK2/STAT5 (Bouscary et al., 2003; Carroll et al., 1991; Ghaffari et al., 2006; Linnekin et al., 1997; Miura et al., 1994; Socolovsky et al., 1999; Wandzioch et al., 2004). In concert with cytokine signaling, several key erythroid-restricted transcription factors (including GATA-1, FOG-1, SCL/TAL-1, EKLF/KLF1) associate with generalized cofactors to activate the transcription of erythroid-specific genes and suppress those of alternate lineages (Akashi et al., 2003; Cantor and Orkin, 2002; Cross and Enver, 1997; Hattangadi et al., 2011; Perkins et al., 1995; Pevny et al., 1991; Shivdasani et al., 1995; Tsang et al., 1997).

While focused studies on erythroid cytokine signaling and transcription factors have generated tremendous functional insights into erythropoiesis, they do not provide a systems-wide view. A comprehensive view of erythroid gene expression has been provided by global analysis of erythroid transcriptomes and the epigenome in purified bulk populations and single cells ((Tusi et al., 2018) and reviewed in (An et al., 2015)). These approaches necessarily use global mRNA levels as proxies of protein abundance and infer signaling activity indirectly. A truly system-wide understanding of post transcriptional and translational mechanisms that drive and coordinate terminal maturation is clearly still lacking. Such a dynamic map would complement transcriptome studies to broadly describe the molecular basis of the pathways involved and to understand how cytokine receptor signaling and transcription factors together shape the erythroid proteome.

In contrast to transcriptome and epigenetic studies of erythropoiesis, relatively few proteome studies to date provide a global analysis of the protein landscape. Due to technical limitations, these studies examined only selected maturation stages in limited depth or focused on defined protein families (Amon et al., 2019; Bell et al., 2013; Chu et al., 2018; Gautier et al., 2018; Gillespie et al., 2020; Liu et al., 2017; Pasini et al., 2006; Roux-Dalvai et al., 2008; Wilson et al., 2016). A recent analysis described dynamic changes in protein expression during *in vitro* erythroid differentiation of CD34^+^ HSPCs (Gautier et al., 2016). However, because relatively large numbers of cells were required for proteomic characterization, this study examined semisynchronous erythroid cultures consisting of cells at different stages of maturation. Given the recent dramatic advances in mass spectrometry and label-free quantitative proteomics (Aebersold and Mann, 2016; Bekker-Jensen et al., 2017), we reasoned that it may now be possible to obtain accurate high coverage proteome and phoshoproteome quantification from relatively low numbers of purified erythroid precursors at distinct developmental stages.

We developed a pipeline combining fluorescence activated cell sorting (FACS) enrichment procedures with our state-of-the-art proteomics workflow. We uncovered the temporal staging of developmental regulation through proteome remodeling. To identify the distinct proteome defining each maturation stage from proerythroblast to orthochromatic erythroblast, we developed a bioinformatic deconvolution approach which revealed stage-specific proteins and protein families. Importantly, our proteomics workflow enabled detection of more than one thousand membrane proteins, and identified distinct combinations of solute carrier (SLC) family proteins as stage-specific maturation markers. Pursuing post-translational regulation further, in-depth sensitive quantitation of the global phosphoproteome with our EasyPhos platform (Humphrey et al., 2015; Humphrey et al., 2018) provided direct evidence for intricate developmental stage-specific regulation by post-translational modification. To functionally explore the identified signaling modules, we performed a kinome-targeting CRISPR/Cas9 screen, which in combination with our proteomic studies, identified distinct signaling requirements for erythroid maturation. Focusing on networks amongst over 27,000 phosphosites and kinase functions uncovered the sequential attenuation of c-Kit and EPOR/JAK2 signaling, pinpointing downregulation of Ras/MAPK signaling in promoting terminal maturation. Our system-wide data provide a wealth of molecular information regarding the functional dynamics of complex phosphosignaling networks in erythropoiesis, expanding our knowledge and data for cellular principles of regulation through proteome remodeling.

## RESULTS

### Establishing stage-specific proteomes of human erythropoiesis

To investigate the remodeling of the proteomics landscape during human erythropoiesis, we cultured human peripheral blood-derived CD34^+^ HSPCs under conditions to support erythroid differentiation (**Methods**). We obtained highly enriched populations of erythroid precursors at specific developmental stages by FACS using CD235a (GYPA), CD49d (ITGA4), and Band 3 (SLC4A1) markers (**Figure 1A-B and Figure S1A**) (Hu et al., 2013). We isolated early maturation stages (progenitors, ProE, EBaso, LBaso) after 7 days of culture, while later maturation stages (LBaso, Poly and Ortho) were purified at day 14. According to published guidelines, LBaso stage precursors were isolated after both 7 days and 14 days of culture using the same markers (**Figure 1B**) Note that SCF was present at 7 days but not at 14 days. Purified cell populations were morphologically homogeneous as judged by May-Grünwald-Giemsa staining (**Figure S1B**). Due to relatively low cell yields, ProE and EBaso populations were combined in equal cell numbers prior to subsequent analysis. The resulting five populations/stages are henceforth color-coded as follows: progenitors (mostly CFU-E) (Hu et al., 2013; Li et al., 2014a; Yan et al., 2017), yellow; ProE/EBaso, blue; LBaso day7, light pink; LBaso day14, dark pink; Poly, dark blue, and Ortho, orange (**Figure 1A-B**).

**Figure 1.**
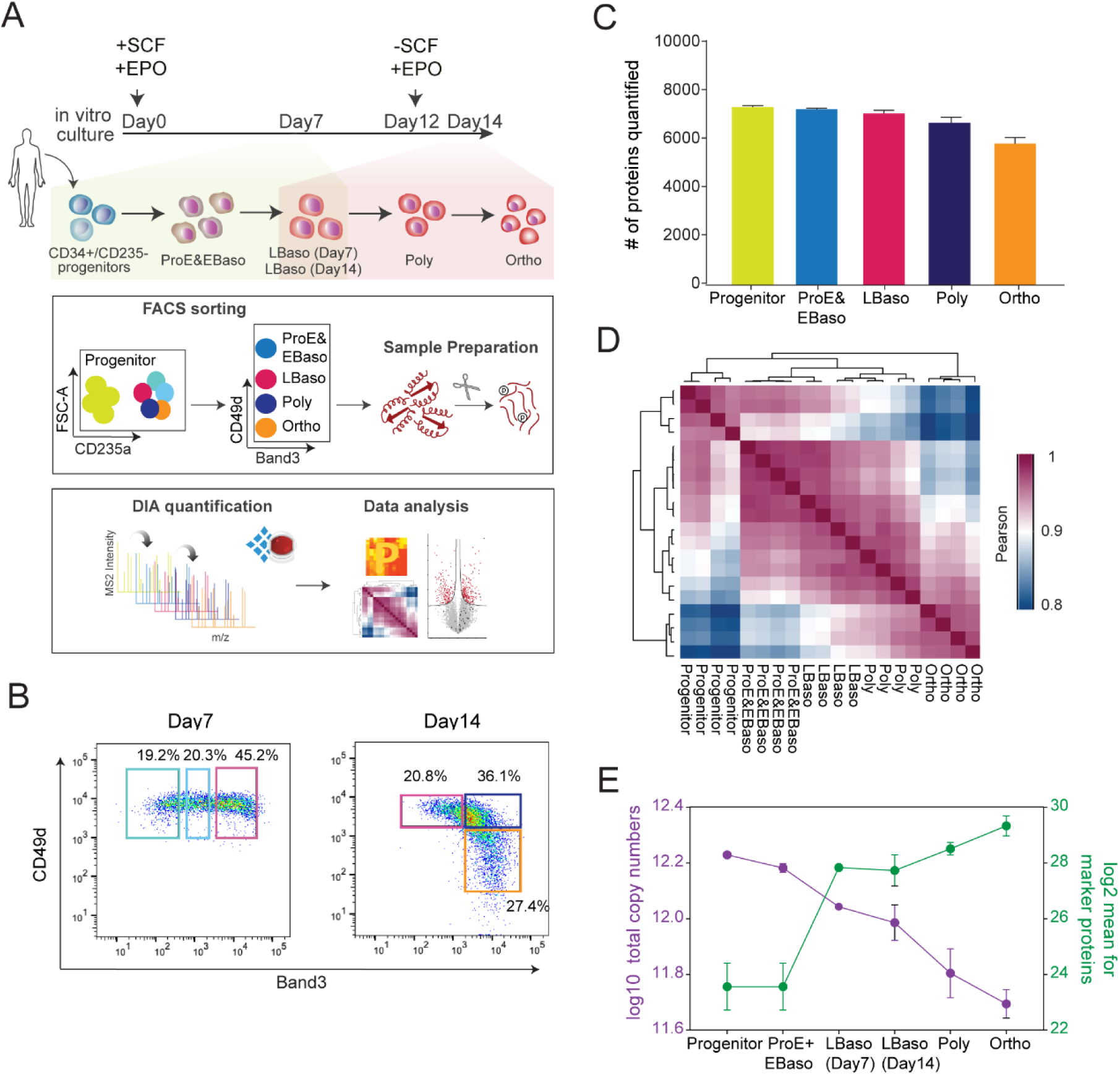
Establishing differentiation stage-specific proteomes of human erythropoiesis. (A) Top panel depicts culture conditions for *in vitro* erythroid differentiation of CD34^+^ cells. Shading indicates the presence of SCF and EPO (yellow), or EPO alone (pink). The lower panels indicat the workflow of our study, including FACS gating/sorting strategy of erythroid precursors and single shot DIA analysis. (B) FACS gating regime to enrich for ProE, EBaso, LBaso, Poly, and Ortho erythroblasts. (C) Number of different proteins quantified in each differentiation stage. (D) Correlation based clustering illustrating the reproducibility between biological replicates. High (1.0) and lower (0.8) Pearson correlations are denoted in pink and blue, respectively. (E) Estimated copy numbers of total molecules (purple) and mean copy numbers of the proteins with GO annotations “erythrocyte maturation” and “heme biosynthesis” (green) per cell across maturation stages.

Each population was processed in four biological replicates and their tryptic peptides were analyzed in single shots in Data Independent Acquisition (DIA) mode **(Methods, Figure 1A)**. To generate a project-specific library, we separated peptides by high pH reversed-phase chromatography into fractions, followed by data-dependent acquisition (DDA) and analysis with Spectronaut. The resultant library contained more than 9,000 protein groups, 7,479 of which could be matched into the DIA runs of at least one maturation stage (q-value less than 1% at protein and precursor levels, **Figure 1C**). In the DIA method, small m/z precursor windows are fragmented in a cyclical manner, which turned out to be crucial for preserving the dynamic range of peptide detection in the presence of the very large hemoglobin peptide peaks that would otherwise complicate analyses at later developmental stages. Remarkably, 84% of all detected proteins were consistently quantified at varying levels across all maturation stages and a relatively small percentage was only matched in a single stage. Quantitative accuracy was high, with Pearson correlations > 0.95 and CVs < 20% for 72% of all proteins between the four biological replicates (**Figure 1D and Figure S1C**). MS signals spanned abundance ranges of five (progenitors) to seven (Ortho) orders of magnitude. As expected, globin proteins increased by approximately one thousand-fold from progenitor to Ortho stage **(Figure S1D)**.

As biological interpretation is facilitated by absolute rather than relative concentration measurements, we employed the ‘proteomic ruler’ method, which uses the fixed relationship between histones and DNA to estimate proteome-wide copy numbers per cell (Wisniewski et al., 2014). Considering that chromatin condensation during erythropoiesis is associated with partial release of major histones from the nucleus and subsequent degradation in the cytoplasm (Zhao et al., 2016a), we first assessed the overall histone content of cells in our system, which indeed declined with progressive differentiation **(Figure S1E)**. Taking this into account for the proteomic ruler calculations, we measured an almost four-fold reduction in total protein copy numbers per cell during differentiation with median copy numbers dropping from 23,380 ± 371 in progenitors to 12,395 ± 1342 at the LBaso stage **(Figure 1E and Table S1)**. In contrast, the average copy numbers of proteins annotated as “erythrocyte maturation” and “heme biosynthesis” by Gene Ontology (GO) increased by approximately 50-fold from progenitor to Ortho stage **(Figure 1E and Table S1)**. Quantitative comparison and copy number estimation of LBaso stages isolated at ether day7 or day14 confirmed their close resemblance at the global proteome level, including marker proteins, such as GYPA, CD49d, Band 3, c-Kit, and several hemoglobin subunits that did not significantly change **(Figure 1E and Figure S2A-C)**. Thus, they were combined for further proteomic analysis unless otherwise noted.

### Dynamic and stage-specific proteome remodeling in erythropoiesis

The five stages of human erythropoiesis clustered separately by principal component analysis (PCA) with very high concordance between replicates **(Figure 2A)**. Hierarchical clustering of 4,316 proteins with statistically different expressions (ANOVA, FDR<0.01), revealed drastic differences in the stage-specific proteomes. Rather than straightforward increase or decrease in protein levels across differentiation, proteins cluster into one of six distinct profiles of temporal co-expression dynamics **(Figure 2B and Table S3)**. In addition to known developmental themes in each cluster, GO enriched terms point to novel state-specific regulation (summarized in **Figure S3**). In pairwise comparisons between successive stages, 2,157 proteins (29%) changed significantly at the first transition **(Figure 2C)**. The overall proteome was more stable from ProE/EBaso to Poly stages, with 8.5% proteins up-or down-regulated. In contrast, almost 20% of the proteome significantly changed in the last investigated transition, reflecting the specialization towards mature erythrocytes **(Figure 2C)**.

**Figure 2.**
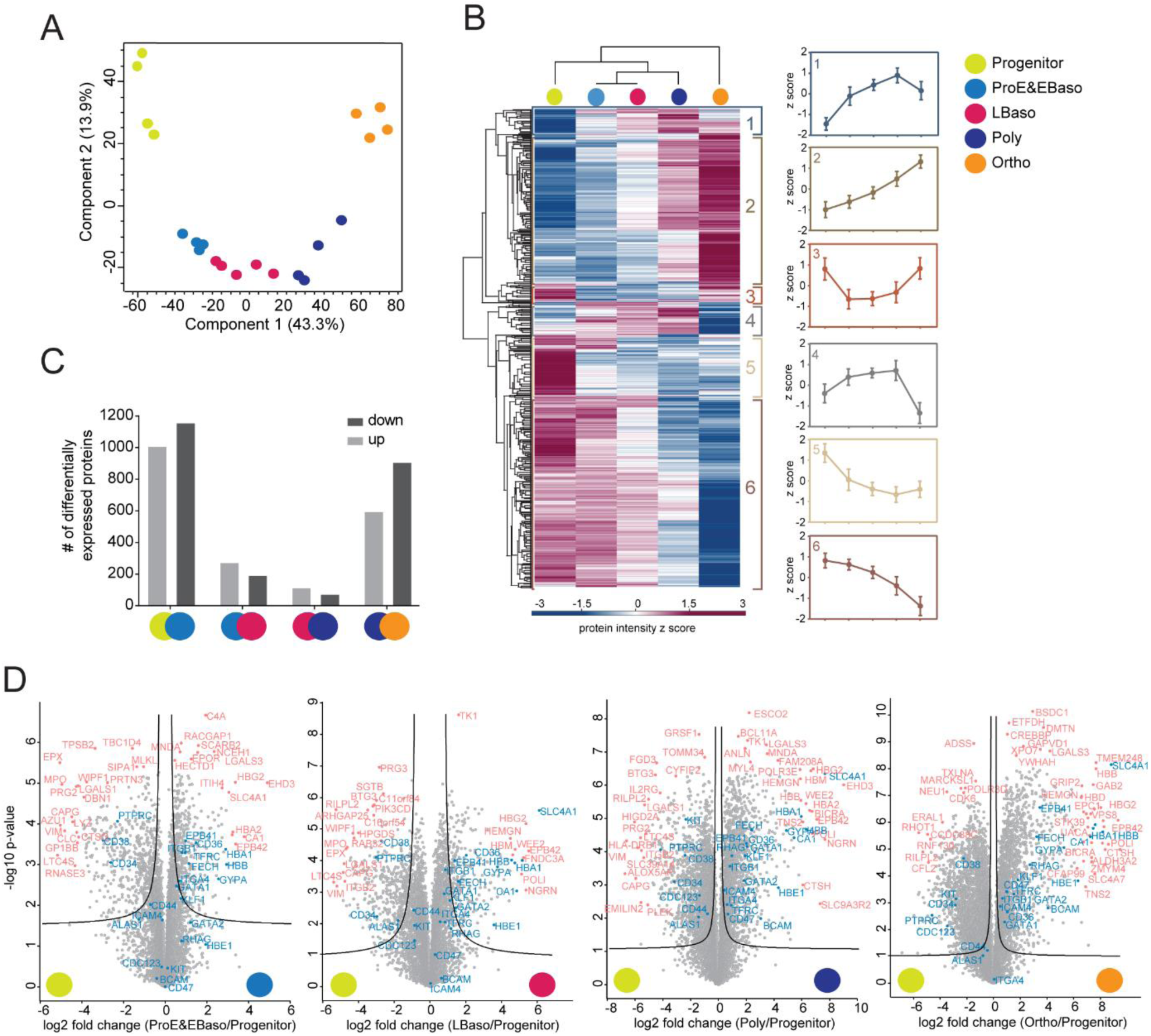
Dynamic and stage-specific proteome remodeling in erythropoiesis. (A) PCA of differentiation stages along with their biological replicates based on their proteomic expression profiles. (B) Heat map of z-scored protein abundances (log2 DIA intensities) of the differentially expressed proteins (ANOVA, FDR<0.01) after hierarchical clustering reveals six main profiles. Mean z-scores with standard errors (SEM) are shown in each stage. (C) Number of differentially expressed proteins in pairwise comparisons of succcessive stages of human erythroid differentiation. (D) Individual Volcano plots of the (-log10) p-values vs. the log2 protein abundance differences between progenitor and the four differentiation stages. Selected significant proteins and previously reported marker proteins are labeled in pink and blue, respectively and significance lines (FDR<0.01) are shown.

To discover unique stage-specific marker proteins we compared all stages against each other **(Figure S4A)**. Interestingly, the Poly stage can be distinguished by the centralspindlin and chromosomal passenger complexes (Benjamini-Hochberg, FDR<0.01). These proteins regulate cytokinesis in the late stages of cell division and also likely participate in erythroblast enucleation. Indeed, mutations in the kinesin KIF23B cause congenital dyserythropoietic anemia associated with erythroid multinuclearity and impaired erythropoiesis (Liljeholm et al., 2013). This analysis, like the ANOVA results, revealed the most drastic proteome changes occurring at the transition from progenitors to ProE/EBaso and from Poly to Ortho **(Figure S4B)**. The cumulative proteome remodeling from the progenitors was reflected in a very large fraction of differentially represented proteins at the later maturation stages, Poly and Ortho (44% and 57%, respectively, two sample test, FDR<0.01 and S0=0.1) **(Figure 2D)**.

Taken together, our stage-specific proteomic data enable accurate, quantitative and in-depth monitoring of global protein expression during human erythropoiesis. The identified proteins are potentially important for the functional specialization of erythroid cells towards mature erythrocytes and represent excellent starting points for more detailed mechanistic studies.

### Dramatic remodeling of the transmembrane proteome in erythropoiesis

Our data captures distinct regulation of proteins that contribute to the highly specialized erythroblast membrane at later developmental stages. Despite identification of several transmembrane proteins as markers of erythropoiesis over the years (Chen et al., 2009), there is still limited systems-wide information on them. Our optimized lysis and digestion protocol enabled unbiased access to the membrane proteome and provided a comprehensive view of membrane proteins during differentiation. Across the differentiation stages, we quantified 1,033 plasma membrane proteins (∼21% of the total genome-encoded plasma membrane proteome in humans and ∼14% in our study), of which 692 changed significantly **(Table S1)**. Our data identifies a plethora of new examples that will aid in pinpointing maturation stages and in better understanding of of erythroid biology.

Of the significantly changing membrane proteins, we could map 86% to pathways, with transport of small molecules across plasma membranes among the most represented (p 8.7 E-09). Further functional classification showed markedly strong enrichment of ‘SLC (solute carrier)-mediated transmembrane transport’ **(Figure 3A)**. The roles of SLCs in biology has arguably been understudied, but now there are systematic efforts characterizing their roles (Cesar-Razquin et al., 2015). Notably, since identification of “Band 3” as a solute carrier protein (SLC4A1) 35 years ago (Kopito and Lodish, 1985), it has become clear that SLC proteins must have widespread roles in erythropoiesis. Remarkably, our data quantified 101 SLCs, 68 of which significantly change in at least one transition **(Figure 3B)**, likely reflecting remarkable changes in metabolic requirements along the stages of maturation. As summarized in **Table S2**, 62 of these have known or purported substrates associated with them.

**Figure 3.**
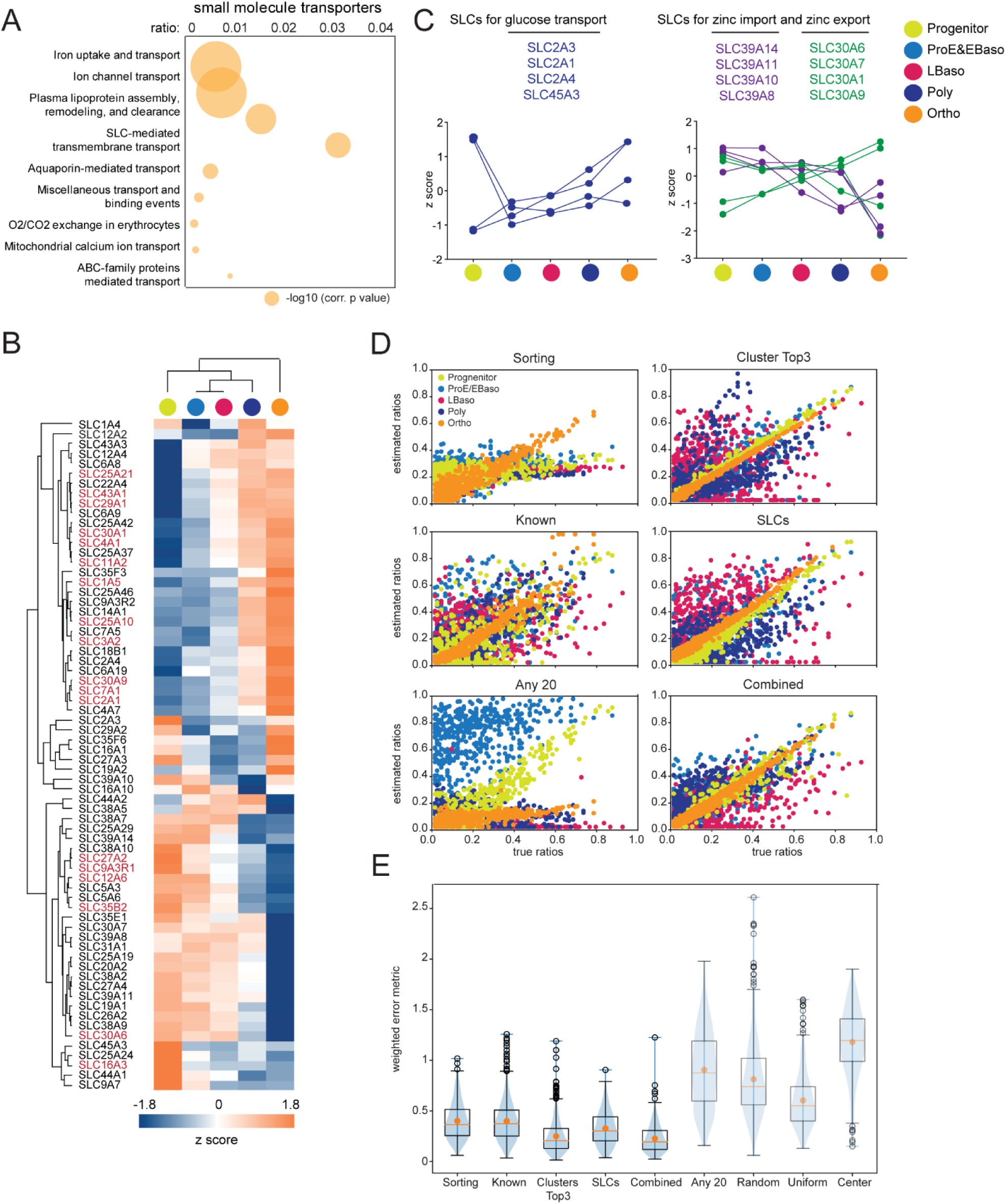
Solute carriers in erythroid maturation and computational extraction of stage-specific protein markers. (A) Plot shows the overrepresented Reactome pathways (Jassal, 2011; Jassal et al., 2020) with their corrected p-values (-log10) and the ratios of given entities from a particular pathway vs all entities from that pathway (n=1,882). (B) Heat map of z-scored SLC protein abundances (log2 DIA intensities) across differentiation. The proteins in red were used to generate the input matrix for the SLCs marker set used in D and E. (C) Expression of four different glucose transporters during erythrocyte development (log2 DIA intensities, left panel). Counterveiling expression regulation of zinc importers and exporters during erythrocyte development (log2 DIA intensities, right panel). (D) Computation sorting quality comparing pre-defined versus estimated ratios of cells in the five differentiation stages. (E) Accuracy of cell type prediction based on different protein marker stets as measured by a weighted error metric (y-axis, also see Methods). The blue violin plots illustrate the underlying distribution reaching from minimum to maximum. The black box plots depict the quartiles of the distribution with whiskers extending to the quartiles ± 1.5 x interquartile range. The orange horizontal lines indicate the median and the orange dot highlights the mean of the distribution.

Only 22 of the significantly regulated SLCs have previously been linked to erythrocytes, erythropoiesis or anemia. For instance, Mitoferrin-1 (SLC25A37), with a continuous upregulation during erythroid maturation, is a mitochondrial iron importer essential for heme biosynthesis in erythroblasts (Shaw et al., 2006). For some SLCs, roles in transporting nutrients including glucose and amino acids, and ions such as zinc, and necessary functions as redox regulators in erythropoiesis have already been described (**Table S2**). In addition, our dataset also contains many transporters – including for vitamins, lipids, and whose substrates have not yet been identified – vastly extending the repertoire of SLCs and transported molecules associated with erythropoiesis.

Among the prominent observations emerging from our data were the several differentially expressed SLCs attributed to a common ligand. We first focused on hexose/glucose transporters. It has been known that cellular metabolism in mature red blood cells is strictly limited to glycolysis, which makes glucose uptake cruical for erythrocyte development. Glucose uptake during maturation appeared to roughly track with EPOR expression, reaching a maximal value when EPO response was highest, perhaps because of regulation by EPO stimulation in erythroid progenitor cells, as reported previously (Rogers et al., 2010). In line with the growing need for glucose during maturation, two out of four identified SLCs transporting glucose (SLC2A1 and SLC2A4) gradually increased from progenitors to Ortho **(Figure 3B-C, Table S2)** and their concordant profiles have been described recently (Justus et al., 2019). The other two glucose transporters (SLC2A3 and SLC45A3) are highly expressed specifically in progenitors, and to our knowledge have not been associated with erythropoiesis; their regulation would be interesting to investigate in the future.

A second remarkable example concerns SLCs for transporting metal ions (14 identified in total) with a full eight of them dedicated to zinc import and export. Maintainence of intracellular zinc levels controlled by GATA/heme circuit has recently been discovered as a vital determinant of erythroid maturation (Tanimura et al., 2018). This indicates the adaptation of differentiating cells to stage-specific metabolic requirements and their interaction with the environment. Apart from the zinc importer SLC30A1 and zinc exporter SLC39A8, previously described in a “zinc switch” model reflecting their reciprocal expression during terminal erythropoiesis (Tanimura et al., 2018), we here uncovered additional three upregulated exporters (SLC39 family) and three downregulated importers (SLC30 family) during maturation **(Figure 3B-C, Table S2)**. Only one of these had prior implications in erythrocyte homeostasis (Ryu et al., 2008), suggesting even more intricate and possibly redundant regulation of zinc homeostasis.

### Computational extraction and characterization of stage-specific protein markers

Given the distinct stage-specific expression patterns of the SLCs, we wondered if they could even serve as marker and selection proteins. The standard approaches for distinguishing erythroid developmental stages rely on canonical cell surface markers, including the ones we employed for FACS enrichments (Chen et al., 2009). Our proteomics analysis revealed that drastic proteome-wide changes of numerous proteins occurred at transitions, in particular from progenitors to ProE/EBaso and from Poly to Ortho, implying the expression of numerous new stage-specific protein markers that might be exploited for refining the isolation and quantification of each differentiation stage. Panels of proteins with characteristic profiles could also be useful for *in silico* deconvolution of mixed developmental populations.

As a starting point, we investigated a *known marker set* of 22 proteins as well as *FACS sorting markers* **(Table S2)**. These proteins correlated well with their expected expression profiles along the differentiation process. We next constituted a marker set of 18 *SLCs* on the basis of the most consistent quantification profiles (**Figure 3B**). As a final set, we selected 18 stage-specific proteins from our proteomics data comprising the top three most significant ones for each of the six clusters in Figure 2A (*cluster Top3 set*, smallest ANOVA q-values) **(Table S2)**. Among those, KLF13, which activates the promoters of several erythroid genes *in vitro*, was gradually upregulated until very late stages, consistent with its reported role in mouse erythroblast maturation (Asano et al., 2000; Gordon et al., 2008).

With these four protein panels in hand (*sorting and known markers, cluster Top3, SLCs*), we developed an *in silico* deconvolution approach to distinguish different developmental stages. Briefly, we used these marker panels to define signature matrices which we applied in 500 random *in silico* mixtures of aggregated abundances from a linear combination of the five differentiation stages at predefined ratios. We evaluated the results by comparing the estimated ratios to the predefined ratios of the *in silico* mixtures **(Figure 3D-E)**.

The *sorting and known markers* reasonably estimated the fraction of the Ortho stage in the mixtures, but performed worse for all other stages (diagonal orange markers in **Figure 3D**). Remarkably, the *cluster Top3 and SLC markers* better characterized the differentiation process than previously known proteins and produced more accurate estimations for both Ortho and progenitor fractions in the computational mixture populations (diagonal orange and yellow markers in **Figure 3D**). However, they were still less effective at distinguishing stages from ProE to Poly, in line with their smaller proteome differences in our data (**Figure 3D**). A combined set of 62 proteins outperformed all others, even in estimating intermediate, adjacent differentiation stages as judged by a quantitative error analysis and compared to random controls **(Methods, Figure 3E)**. In addition to recent advances in single cell transcriptomics (Tusi et al., 2018), our deconvolution approach could further aid the identification of specific populations amongst bulk pools obtained during erythropoiesis, for example in the study of differentiation dynamics from *in vivo* samples. The proteins selected in this analysis, especially the *SLCs* add to our resource as they are interesting candidates for investigating stage-specific mechanisms in follow up studies.

### An orchestrated network of erythropoietic kinases and their downstream targets

Several kinases act in or have already been implicated in a complex regulatory network in erythropoiesis. To advance our understanding of the dynamic phospho-regulatory network during erythropoiesis, we assessed temporal kinase activities at a global scale across terminal maturation. Mining of our proteome data revealed an astounding 270 kinases and 90 phosphatases that were differentially expressed with clear stage-specific profiles during differentiation **(Figure 4A)**. To investigate their activities, we turned to phosphoproteomics which globally captures their substrates **(Figure 4B)**. We enriched phosphopeptides from the same differentiation stages in biological quadruplicates using the EasyPhos platform **(Figure 4B)** (Humphrey et al., 2015; Humphrey et al., 2018). This streamlined protocol enabled deep profiling of phosphoproteomes at specific developmental stages in single-run DDA measurements from only 80 µg of protein lysates, capturing 27,166 distinct phosphosites on more than 4,200 proteins **(Figure 4B)**. Almost 20,000 sites were identified in more than two replicates of at least one maturation stage and 3,604 were novel sites according to the PhosphoSitePlus database (Hornbeck et al., 2012) **(Figure 4C and Table S3)**. Given the prominent changes in the plasma membrane proteome, it was interesting to see that 401 of them had phosphosites, often multiple ones within proximity in linear sequence stretches. This encompassed 23 of the aforementioned SLCs, sugesting stage-specific signaling roles (Taylor, 2009) in addition to their dynamic expression across stages. Specifically, our phosphoproteomics also identified Ser/Thr phosphorylation sites on Band 3, whose tyrosine phosphorylation is known to enable docking of cytoplasmic signaling molecules (Brunati et al., 2000; Yannoukakos et al., 1991).

**Figure 4.**
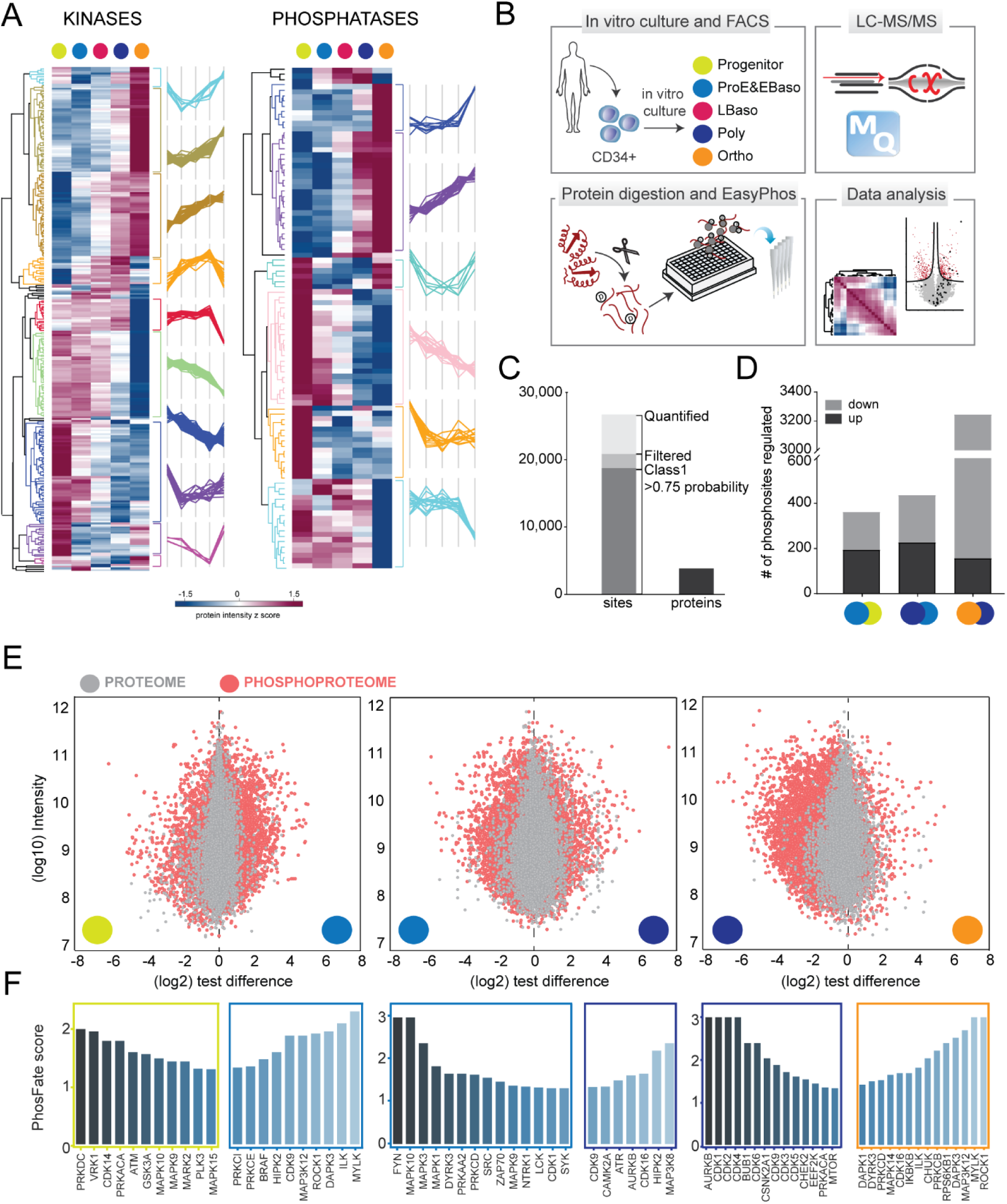
An orchestrated network of erythropoietic kinases and their downstreat targets. (A) Heat map of z-scored and differentially regulated kinase and phosphatase abundances (log2 DIA intensities) across differentiation. (B) Experimental design of the phosphoproteomic study, performed on the same populations as collected for the full proteome analyses (also see Figure 1A). Analytical workflow including phospho-enrichment, single shot DDA acquisition and data analysis. (C) Number of identified and quantified Class 1 sites (localization probability to a single amino acid > 0.75) after filtering for 50% data completeness in at least one differentiation stage. Total number of phosphoproteins is also shown. (D) Significantly regulated phosphorylated sites in pairwise comparisons of ProE/EBaso vs Progenitor, Poly vs ProE/EBaso, and Ortho vs Poly. (E) Distributions of phosphopeptides and their matching proteins based on their log10 intensities (Y-axis) vs log2 test differences (X-axis) are illustrated for ProE/EBaso vs Progenitor (left), Poly vs ProE/EBaso (middle), and Ortho vs Poly (right). Pink represents phosphopeptides whereas grey represents proteins. (F) Stage-specific predicted active kinases based on targeted sites identified by PhosFate profiler (http://phosfate.com). Left boxes represent kinases whose substrates are preferentially detected at the earlier stage of differentiation and right boxes represent those whose substrates are preferentially detected at the later stage.

For further statistical analysis, we used a stringently filtered dataset of 12,216 phosphopeptides quantified in all four replicates of at least one differentiation stage. Strikingly, about half of these phosphosites significantly changed in at least one developmental transition (ANOVA, FDR<0.05) and a quarter of all phosphosites (3,089) were dephosphorylated from Poly to Ortho stage **(Figure 4D and Table S3)**.

To compare the dynamics of the phosphoproteomes to the proteomes, we visualized fold change distributions of quantified proteins (grey) and phosphopeptides (pink) for three pairwise comparisons: (i) progenitor vs ProE/EBaso, (ii) ProE/EBaso vs Poly, and (iii) Poly vs Ortho (**Figure 4E**). The fold change distributions of phosphopeptides were considerably more scattered than those of proteins in all three comparisons, reflecting dynamic, large scale phosphoregulation. The largest fold change of regulated phosphopeptides occurred between early stages of progenitor to ProE/EBaso and the later stages, Poly to Ortho (**Figure 4E**). The highly dynamic changes in global phosphorylation landscape likely reflects critical roles for distinct kinases at specific maturation stages.

Next, we inferred kinase activities from the phosphoproteome by stage-dependent enrichment analysis using PhosFate profiler (**Figure 4F**) (Ochoa et al., 2016). This method predicts changes in kinase activity by testing the enrichment of differentially regulated, annotated kinase-substrate motifs. Substrates peaking during the early stages of differentiation (ProE/EBaso) were enriched with motifs for kinases of the MAPK signaling network (BRAF, MAPK1, MAPK3, FYN, SRC), which are known to promote cell cycle and proliferation (Carroll et al., 1991; Geest and Coffer, 2009; Sakamoto et al., 2000). Interestingly, the observed substrate phosphorylations suggest that CDK1 and many other cell cycle associated kinases (AURKB, BUB1, CDK14, CDK16, CDK2, CDK3, CDK4, CDK5, CDK6, and DYRK3) remain active until very late stages (Poly and Ortho). DNA damage checkpoint kinases (ATM, ATR, and CHEK2) were also enriched, presumably to maintain genome stability during erythroid differentiation. Together, our data reveal a rich network of temporally activated kinases during differentiation of human erythrocytes.

### CRISPR/Cas9 screen reveals critical functions of the erythropoietic kinome

The proteomics analysis established a “kinome atlas” revealing dynamic changes in kinase abundance and activity at distinct stages of erythropoiesis, with a dramatic decrease in the global phosphoproteome during late maturation. To investigate potential functional implications of these changes, we performed a CRISPR/Cas9 screen in HUDEP-2 cells, an immortalized human erythroblast line that proliferates in an immature state and can be induced to undergo terminal maturation by manipulation of culture conditions (Kurita et al., 2013). HUDEP-2 cells stably expressing Cas9 (HUDEP-2^Cas9^) were transduced at low multiplicity of infection with a lentiviral vector library encoding 3,051 single guide (sg) RNAs targeting the coding regions of most known kinases (n=482) and a green fluorescence protein (GFP) reporter gene (Grevet et al., 2018; Tarumoto et al., 2018) (**Figure 5A**). Two days later, GFP^+^ cells were flow cytometry-purified, and split into pools for further expansion or induced maturation, followed by next generation sequencing (NGS) to assess sgRNA abundance **(Figure 5A and S5A-D)**. Compared to cells at two days post-transduction (“day0”), 30 sgRNAs were underrepresented after 10 days of expansion (FDR<0.05), reflecting candidate kinase genes that promote survival and/or proliferation of immature erythroblasts **(Figure 5B and Table S4)**. These genes encoded cyclin-dependent kinases (*CDK1, CDK7, CDK9*), cell signaling components (*KIT, JAK2*), DNA damage checkpoint response proteins (*ATR, CHEK1*) and a regulator of ion flux (*OXSR1*), several of which exhibited maturation stage-specific expression in our proteome analysis (e.g. *CDK1, CDK9, ATR, KIT*, and *JAK2*) **(Figure 4F)**. The *KIT* and *JAK2* genes are essential signaling molecules for erythropoiesis (Munugalavadla and Kapur, 2005; Neubauer et al., 1998; Parganas et al., 1998). Previous proteomic studies identified OXSR1 (OSR, oxidative stress-responsive kinase 1) as one of the most abundant Ser/Thr kinases in reticulocytes and mature erythrocytes (Gautier et al., 2018). The OXSR1 protein phosphorylates Na^+^–K^+^ and K^+^–Cl^−^ membrane co-transporters to activate and inhibit their activities, respectively (de Los Heros et al., 2014). Our data suggest a role for OSXR1 in the maintenance of erythroid precursors.

**Figure 5.**
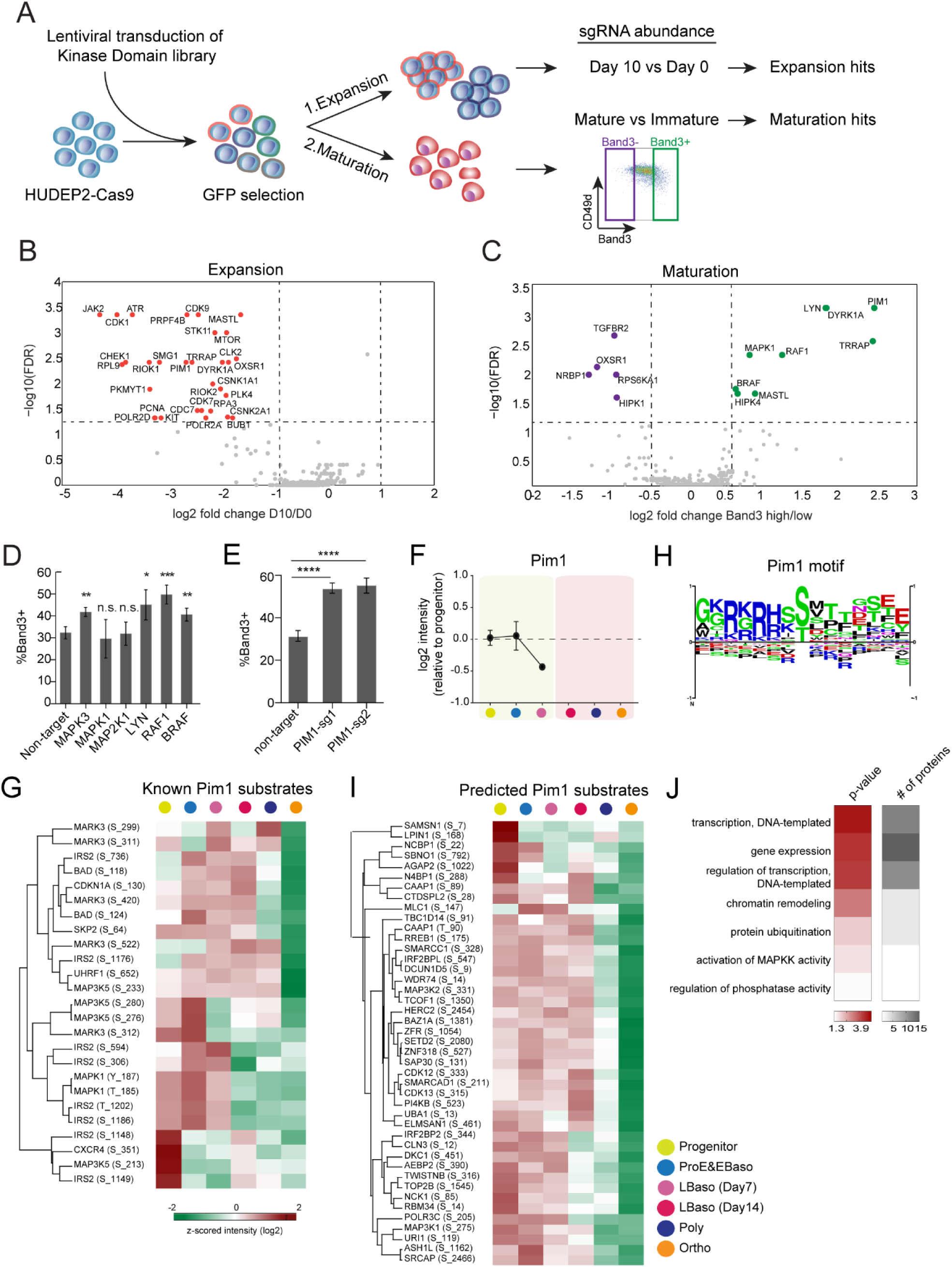
Kinome-targeting CRISPR/Cas9 screen in HUDEP-2 cells. (A) Workflow of a CRISPR/Cas9 screen with an sgRNA library targeting 482 human kinase genes to identify those that alter erythroid precuresor expansion or terminal maturation. (B) Volcano plots showing FDR vs log2 fold-change in sgRNA abundance between Day 0 and Day 10 of expansion. Results were analyzed using Mageck (Methods). Each dot represents a single kinase gene based on the enrichment of four sgRNAs. Significantly different genes (FDR<0.05; log2fold-change<-1) are shown in red. (C) Volcano plots showing FDR vs log2 fold-change in sgRNA abundance between Band3 high and low fractions after three days of maturation. Results were analyzed using Mageck (Methods). Each dot represents a single kinase gene based on the enrichment of four sgRNAs. The significant positive or negative regulators for maturation (FDR <0.05; log2 fold-change<-0.5 and log2 fold-change>0.5) are shown in purple and green, respectively. (D) HUDEP-2 cells expressing Cas9 were were transduced with lentiviral vectors encoding single sgRNAs targeting the indicated genes, induced to undergo terminal maturation and analyzed after 3 days. Graph shows fraction of Band3^+^ cells. Error bars represent mean ± SEM of 3 biological replicates. \**P* < 0.05, \*\**P* < 0.01, \*\*\**P* < 0.005; n.s., not significant; unpaired t-test. (E) Effects of two different PIM1-targetign sgRNAson erythroid maturation of HUDEP-2 cells, performed as described for panel D. Error bars represent mean ± SEM of 3 technical replicates. \*\*\**P* < 0.005, \*\*\*\**P* < 0.0001; unpaired t-test. (F) Protein abundances (log2) of PIM1 during terminal differentiation of primary erythroblasts. Yellow highlighting indicates that SCF and EPO were present in the culture medium, while pink indicates EPO only. (G) Heat map showing z-scored (log2) phosphopeptide intensities detected in known PIM1 substrates. (H) Consensus PIM1 kinase motif from PhosphoSitePlus database (Hornbeck et al., 2012). (I) Heat map of z-scored (log2) phosphopeptide intensities of potential PIM1 kinase targets identified by motif analysis. (I) Gene Ontology (GO) enrichment analysis of potential PIM1 kinase targets, performed using Fischer’s exact test. 5% threshold was applied to Benjamini-Hochberg FDR to determine the significance.

Transduced HUDEP-2 cells induced to undergo terminal maturation were cultured for 3 days, fractionated according to their expression of the late-stage erythroid marker Band 3, and analyzed by NGS for sgRNA abundance. Single guide RNAs for five genes were significantly overrepresented in immature (Band3^-^) cells, indicating that these genes are positive effectors of maturation, while sgRNAs for nine genes were overrepresented in mature (Band3^+^) cells, representing candidates that inhibit maturation **(Figure 5C and Table S5)**. There was minimal overlap between genes that affect expansion or maturation **(Figure S5E)**. Notably, eight kinases identified as regulating differentiation in the CRISPR/Cas9 screen are also identified amongst the stage-specific active kinases predicted by phosphorylation of their cognate motifs **(Figure 4F and Figure S5F)**.

We noted that disruption of numerous genes stimulating the Ras/MAPK signaling pathway caused accelerated erythroid maturation (**Figure 5C**). Three of the corresponding proteins, RAF1, BRAF1 and MAPK1, are members of the canonical Ras/MAPK family, while LYN is known to engage and potentiate RAF1 (Tilbrook et al., 2001). In non-erythroid cells, PIM1 kinase has been shown to phosphorylate ERK and activate Ras/MAPK signaling (Wang et al., 2012). To validate these candidates, we transduced Cas9-expressing HUDEP-2 cells with individual sgRNAs for each gene and then induced erythroid maturation. Consistent with results of the screen, knockout of *RAF1, BRAF1, MAPK3, LYN* or *PIM1* resulted in significantly accelerated terminal maturation **(Figure 5D-E and S5G-I**). Together, these findings indicate that downregulation of the Ras/MAPK pathway promotes terminal erythroid maturation. Consistent with this hypothesis, the *TGFRB2* gene, identified as positive regulator of maturation (**Figure 5C**), is known to inhibit MAPK signaling (Li et al., 2014b).

Results of the screen identified PIM1 as a candidate gene that both drives erythroid precursor expansion and inhibits maturation (**Figures 5B-C and S5H**). Consistent with the latter, PIM1 protein levels were undetectable after the LBaso stage (**Figure 5F**). The *PIM1* gene encodes a serine-threonine kinase oncoprotein that stimulates cell survival and cell cycle progression by phosphorylating numerous substrates that have been identified in non-erythroid cell types. To identify potential effectors of PIM1 signaling in erythroid cells, we searched our phosphoproteomics data for previously-described PIM1 substrates (Hornbeck et al., 2012) and identified 25 significant phosphosites (out of 100) that coincided with PIM1 expression **(Figure 5G)**. We then generated a PIM1 phosphorylation site consensus motif using the PhophoSitePlus database (Hornbeck et al., 2012) **(Figure 5H)**, and investigated whether the motif is enriched in the erythroid maturation stage-dependent phosphoproteome. This identified 79 phosphorylation targets of which 54% have maturation stage-dependent profiles correlating with PIM1 protein abundance **(Figure 5I)**. GO-term analysis revealed significant enrichment of terms associated with “chromatin remodeling”, “transcriptional regulation”, “kinase/phosphatase activity”, and “ubiquitylation” (**Figure 5J**). Of particular interest were Ras/MAPK family members such as MAP3K1, MAP3K5 and MAP3K2, consistent with a regulatory role of PIM1 in Ras/MAPK signaling.

### System-wide dissection of c-Kit and EPOR phosphosignaling in erythropoiesis

Phosphoprotein analysis and a CRISPR-Cas9-sgRNA screen defined a dynamic, developmental stage-specific kinome during erythropoiesis and indicated that Ras/MAPK downregulation might be critical for erythroid maturation. We explored this further by examining our proteomics dataset for Ras/MAPK signaling components in relation to the expression and activities of c-Kit and EPOR. The kinetics of Ras/MAPK protein expression varied across erythroid maturation with MAPK1, MAPK3 and RAF1 persisting into late maturation stages **(Figure 6A)**, suggesting that their activity may be suppressed post-translationally. In agreement, activating T185/Y187 phosphorylations on ERK indicated maximal activity during ProE/EBaso and termination by the LBaso stage **(Figure 6B)** (Gupta and Prywes, 2002; Michaud et al., 1995). The activating S63 phosphorylation on ATF1, a distal target of Ras/MAPK signaling, peaked later (at the LBaso stage) and persisted throughout erythroid maturation **(Figure 6B)**. The RSK kinase, which phosphorylates ATF1, is activated by both MAPK and PI3K/Akt-mTOR signaling (Koh et al., 1999).

**Figure 6.**
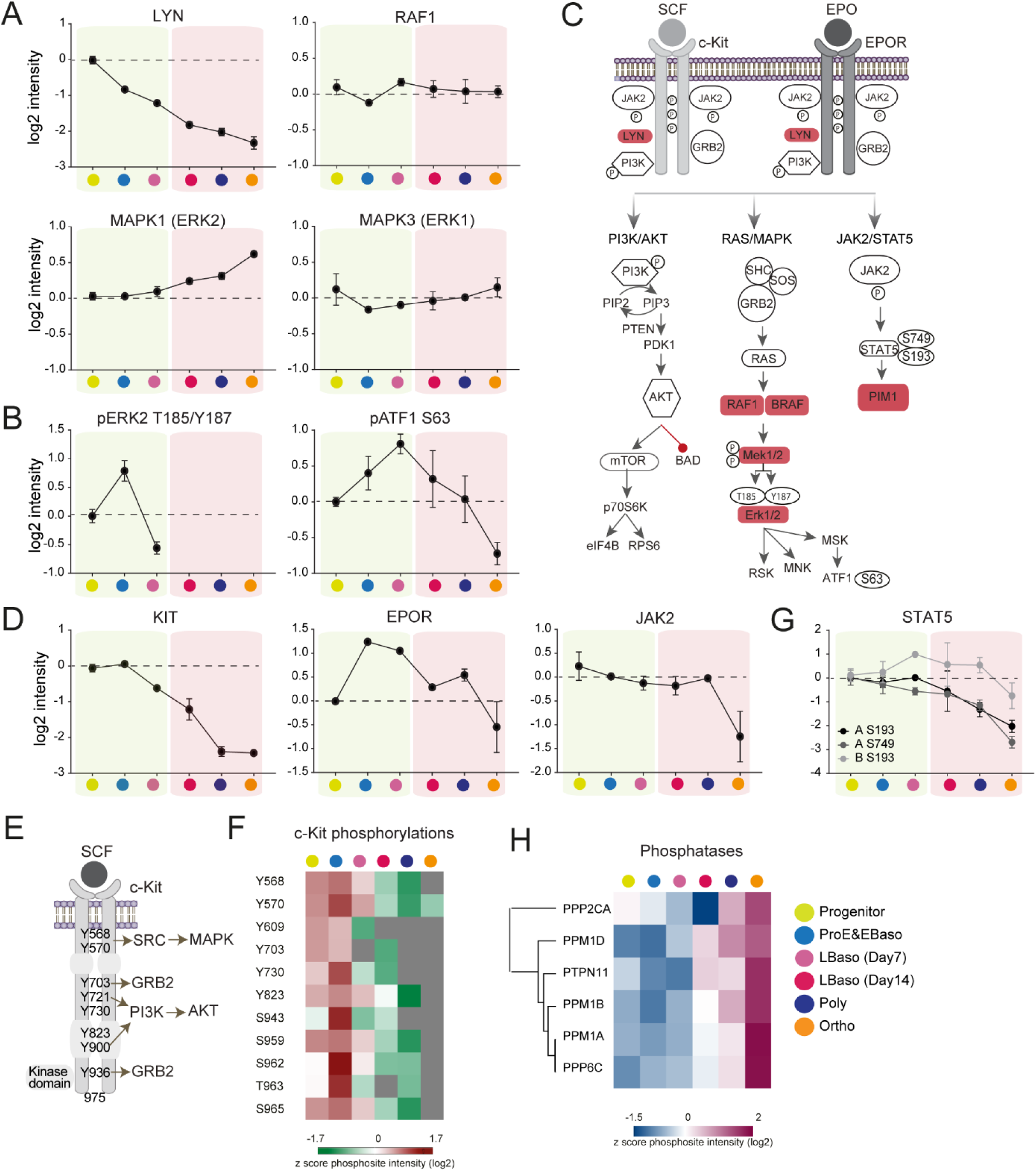
System-wide dissection of c-Kit and EPOR phosphosignaling. (A) Protein abundances (log2 DIA intensities) normalized to progenitor stage. In all panels, stages shaded yellow were cultured with SCF and EPO, while those shaded pink were cultured with EPO only. Data is plotted if quantified in at least 50% of biological replicates. Error bars represent mean ± SEM of at least two biological replicates. (B) Profiles of phosphorylations (log2 DDA intensities) normalized to progenitor stage. (C) Major signaling pathways downstream of c-Kit and EPOR activation by their corresponding ligands stem cell factor (SCF) and erythropoietin (EPO). Shaded genes indicate those indentified to inhibit erythroid maturation in the CRISPR/Cas9 screen described in Figure 5. (D) Protein abundances (log2 DIA intensities) of c-Kit, EPOR, and JAK2, normalized to progenitor stage. (E) Following activation by SCF, phosphorylated tyrosine residues on c-Kit receptor serve as binding sites to key signal transduction molecules (SRC, GRB2, and PI3K) resulting in activation of downstream signaling pathways. (F) Heat map of z-scored (log2) phosphopeptide intensities of c-Kit receptor. (G) Profiles of STAT5A/B phosphorylations (log2 DDA intensities) normalized to progenitor stage. (H) Heat map of z-scored protein abundances (log2 DIA intensities) of phosphatases.

The erythroid cytokine receptors c-Kit and EPOR have distinct roles in erythropoiesis, although their signaling pathways overlap considerably **(Figure 6C)**. Our erythroid culture system contained both SCF (c-Kit Ligand) and EPO in the first and second phase (day0-7) and EPO only in the third phase (day12-14) **(Figure 1A)**. The rationale for this culture system is based on findings that persistently elevated SCF-c-Kit signaling inhibits terminal erythroid maturation (Haas et al., 2015; Munugalavadla et al., 2005; Muta et al., 1995). The levels of c-Kit and EPOR/JAK2 proteins decreased during differentiation but with differing kinetics **(Figures 6D)**. c-Kit levels declined after the ProE stage, similar to ERK activity. Tyrosine phosphorylation in the c-Kit cytoplasmic domain, which reflects receptor activity (Lennartsson and Ronnstrand, 2012), was maximal in erythroid progenitors and ProE/EBaso and decreased by the LBaso stage, even with SCF present in the culture media (**Figure 6D**). ERK2 phosphorylation declined at the same stage, even with SCF present in the culture medium (LBaso day7) and was undetectable in LBaso day14 when SCF was not present in the medium. Thus, c-Kit protein levels and its phosphorylation, along with downstream Ras/MAPK signaling are downregulated relatively early in erythroid maturation, consistent with the literature (**Figure 6E-F**) (Gautier et al., 2016; Matsuzaki et al., 2000).

Compared to c-Kit, EPOR/JAK2 levels were stable until the Poly stage. We were not able to detect phosphorylation of EPOR or JAK2, perhaps because the levels of these proteins are relatively low. Compared to ERK, STAT5 phosphorylation declined more slowly and persisted until the later stages of erythropoiesis, similar to the kinetics of EPOR expression (**Figure 6G**). Thus, ERK phosphorylation levels parallel the expression and activation of c-Kit, while phospho-STAT5 levels correlate with expression of EPOR, likely reflecting preferential signaling activities of the two cytokine receptor pathways. Together, our findings suggest that Ras/MAPK activity occurs during early stages of erythropoiesis, delays terminal maturation and is c-Kit driven. This is consistent with the established role for c-Kit in supporting proliferation and survival of early erythroid progenitor cells and a requirement for c-Kit downregulation during normal erythropoiesis (Bernstein et al., 1991; Muta et al., 1995; Nocka et al., 1989).

Phosphorylation is also regulated by phosphatases. Of 51 phosphatases implicated in inhibiting the Ras/MAPK pathway (Kondoh and Nishida, 2007; Li et al., 2013), 16 were detected in our data, 6 of which were induced during terminal maturation (**Figure 6H**). The majority of these phosphatases are novel candidate genes whose roles in erythropoiesis need further exploration.

## DISCUSSION

Here we show that in-depth quantitative proteomic and phosphoproteomic analysis of purified erythroid precursors at distinct maturation stages is now made possible by state-of-the art MS-based proteomics and our EasyPhos technology, allowing us to assess erythroid maturation at the level of the proteins, the main functional cellular entities. The breadth and depth of coverage achieved by these technologies offers unbiased system wide insights into the regulation of erythropoiesis, which we complemented further by performing an unbiased CRISPR/Cas9 screen to interrogate the erythroid kinome.

Our analyses of proteins mediating solute transport and phospho-based signaling highlight two examples by which the data can be mined for hypothesis generating discovery and focused problems related to erythrod biology. The key emerging concept is that proteome-wide changes accompanying differentiation from early erythroid progenitors into nearly mature erythrocytes involves remarkable regulation of and by signaling pathways – at the protein level.

Tracking the levels across particular families of proteins defined numerous distinctive stage-specific profiles, exemplified by coordinated expression of specific cohorts of SLCs, kinases, and phosphatases. For some of these SLCs, previous reports established roles in transporting crucial molecules for erythropoiesis (**Table S3)**. However, the much larger repertoire of SLCs of unknown biological function likely reflects erythroid stage-specific metabolic requirements to be elucidated in future studies. Beyond this, both the substrate diversity of SLCs and their widespread phosphorylation (average of three per SLC) point toward exquisite fine-tuning and coordination of transporters with stage-specific signaling pathways. So far, very few examples of this mode of regulation have been described, most prominently the tyrosine phosphorylation of the SLC Band 3, which mediates docking of cytoplasmic signaling molecules (Brunati et al., 2000). The prevalence of stage-specific SLC phosphorylation observed here may indicate system-wide coordination of small molecule transport with cytoplasmic signaling throughout erythroid maturation and across many transporters. SLCs are relevant to therapeutics of several human diseases and to drug discovery, either as drug targets themselves or as mediators of drug uptake (Cesar-Razquin et al., 2015). It now seems likely that the varying expression of SLCs with overlapping selectivities, as well as their regulation by post-translational modification, will also contribute to pathology and provide opportunities for therapeutic development (Noomuna et al., 2020). Importantly, our data provide a framework for systemwide studies of SLC small molecule flux and signaling throughout the differentiation process. The dynamics of SLCs, together with more than 700 other quantified membrane proteins may furthermore contribute to our understanding of changing cell membrane properties required for erythropoiesis.

The distinct cohorts of kinases and phosphatases expressed coordinately and with varying kinetics across the erythrocyte maturation pathway likewise reflect extensive protein-level regulation, in this case through post-translational modification. With this notion in mind, we complemented the quantitative stage-specific proteome measurements with a kinome-targeting CRISPR/Cas9 screen and phosphoproteomics, which provides a profile of system-wide signaling across erythroid maturation. Pursuing PIM1, the highest scoring hit for erythroid maturation of HUDEP-2 cells, we defined a composite profile based on (1) stage-specific expression, (2) phosphorylation kinetics of known substrates, and (3) a PIM1 consensus sequence by correlation. Further analysis then provided a list of candidate PIM1 substrates that kinetically parallel PIM1 activity and may coordinate PIM1 activity with that of other diverse signaling effectors including epigenetic regulators, regulators of ubiquitin signaling, and more, whose functional roles in erythropoiesis can now be studied.

We also took advantage of our data to mine the stage-resolved phosphoproteomics of the SCF- and EPO-triggered signaling network, based on its crucial role in erythropoiesis, and opportunities offered by our culturing and FACS-based protocol. Combining phosphoproteomics and a functional screen, our data highlights a general decrease in kinase activity across the erythroid proteome during terminal maturation and in particular a critical role for downregulation of Ras/MAP kinases activity is suggested. Previous studies have demonstrated defective terminal maturation in systems expressing either constitutively active forms of c-Kit or Ras proteins (Haas et al., 2015; Matsuzaki et al., 2000). Our systems-wide analysis of erythropoiesis derived from primary healthy donor human CD34^+^ cells suggests c-Kit drives expansion of early erythroid precursors and inhibits terminal maturation via Ras/MAPK signaling whereas EPOR drives signaling predominantly via the JAK-STAT5 pathway to foster later stages of terminal maturation. These examples demonstrate the ability to gain insight on regulation of complex signaling systems during erythropoiesis using our phosphoproteomics dataset and provide a framework for future studies to interrogate stage-specific erythroid cytokine signaling and regulatory pathways that dampen this signaling.

Given the unexpectedly large role of phosphosignaling during erythropoiesis defined by our unbiased global, high resolution proteomic study, it will now be interesting to investigate other post translational protein modifications, which could employ different enrichment steps but similar strategies for bioformatic and functional follow up. In this regard, we already observed distinct regulation of more than a hundred members of the ubiquitin machinery, making this post-translational modification particularly exciting for further explorations.

## ACKNOWLEDGEMENT

This work was supported by the Max-Planck Society for the Advancement of Science and by the Deutsche Forschungsgemeinschaft (DFG, German Research Foundation) – SCHU 3196/1-1”. We thank Florian Meier, Igor Paron, Christian Deiml, Philipp Geyer, Johannes B Mueller, Fynn M Hansen, Sebastian Virreira Winter and all the members of the departments of Proteomics and Signal Transduction and Molecular Machines and Signaling at Max-Planck-Institute of Biochemistry for their assistances and helpful discussions. We also thank the NCI Cancer Center grant to St. Jude (NIHP30CA021765) and St Jude Core facilities including Flow Core for cell sorting, Hartwell Center for NGS, and Image Core for Cytospin Scanning.

## AUTHOR CONTRIBUTIONS

OK performed proteomics experiments and analyzed the data. PX and YY performed FACS sorting, tissue culture experiments and CRISPR/Cas9 screen and biological validation assay for PIM1. IB developed the bioinformatics deconvolution approach to validate markers. ARFC and ASD helped with the bioinformatics analysis of phosphoproteome data. SVB helped with the analysis and the interpretation of the data. OK, PX, AFA, SVB, BAS, MW and MM designed the study and wrote the paper. BAS, AFA, MW and MM coordinated and supervised.

## DECLARATION OF INTERESTS

The authors declare no competing interests.

## STAR METHODS

### CD34^+^ cell culture and manipulation

Human CD34+ cells were obtained under human subject research protocols that were approved by local ethical committees: St. Jude Children’s Research Hospital protocol “Bone marrow for hemoglobinopathy research” (NCT00669305). CD34+ hematopoietic stem and progenitor cells (HSPCs) were mobilized from normal subjects by granulocyte colony-stimulating factor, collected by apheresis, and enriched by immunomagnetic bead selection using an autoMACS Pro Separator (Miltenyi Biotec), according to the manufacturer’s protocol. At least 95% purity was achieved, as assessed by flow cytometry using a PE-conjugated anti-human CD34 antibody (Miltenyi Biotec, clone AC136, #130-081-002). A 3-phase culture protocol was used to promote erythroid differentiation and maturation. In phase 1 (days0–7), cells were cultured at a density of 10^5^–10^6^ cells/mL in IMDM with 2% human AB plasma, 3% human AB serum, 1% penicillin/streptomycin, 3 IU/mL heparin, 10 µg/mL insulin, 200 µg/mL holo-transferrin, 1 IU EPO, 10 ng/mL SCF, and 1 ng/mL IL-3. In phase 2 (days8–12), IL-3 was omitted from the medium. In phase 3 (days12–18), cells were cultured at a density of 10^6^/mL, with both IL-3 and SCF being omitted from the medium and the holo-transferrin concentration increased to 1 mg/ml. Erythroid differentiation and maturation were monitored by flow cytometry, using FITC-conjugated anti-CD235a (BD Biosciences, clone GA-R2, #561017), APC-conjugated anti-Band3 (gift from Xiuli An Lab in New York Blood Center), and VioBlue-conjugated anti-CD49d (Miltenyi, clone MZ18-24A9, #130-099-680).

### CRISPR/Cas9 screen with kinase-domain library

The kinase domain-focused sgRNA library was designed based on the human kinase gene list from a previous study (Manning et al., 2002). The kinase enzymatic domain information was retrieved from NCBI database conserved domain annotation. Six independent sgRNAs were designed for targeting each individual domain regions. All the sgRNAs were designed using the same design principle reported previously and the sgRNAs with the prediction of high off-target effect were excluded (Hsu et al., 2013). Domain targeting and positive/negative control sgRNAs were synthesized in duplicate or triplicate in a pooled format on an array platform (Twist Bioscience) and then PCR cloned into the BsmB1-digested LRG2.1 vector (Addgene: #108098) using Gibson Assembly kit (NEB). Approximately 12 × 106 HUDEP-2 cells stably expressing Cas9 were transduced at a multiplicity of infection (MOI) of ∼0.3 to minimize the transduction of any cell with more than 1 vector particle and achieve an approximately 1000-fold library coverage (100ul Virus per 2 M cells) such that 40% cell were GFP positive. Two days after infection, GFP^+^ cells were sorted by FACS and then maintained in the expansion medium for 6 days (total 8 days post-infection). Then, half of total cells were kept in expansion culture for additional 10 days and the other half were transitioned to differentiation media and induced maturation for 3 days. Erythroid maturation was monitored by flow cytometry, using FITC-conjugated anti-CD235a (BD Biosciences, clone GA-R2), APC-conjugated anti-Band3 (gift from Xiuli An Lab in New York Blood Center), and Violet Blue–conjugated anti-CD49d (Miltenyi, clone MZ18-24A9). Band3^+^ and Band3^−^ cell populations from the CD235a^+^ cell fraction was purified by fluorescence-activated cell sorting (FACS). Library preparation and deep sequencing were performed as previously described 46,47. Briefly, genomic DNA was extracted using the DNeasy Blood and Tissue kit (Qiagen). Reactions were done with 24 cycles of amplification with 200 ng of gDNA in 25 µL CloneAmp enzyme system and 8 parallel reactions were performed to maintain sgRNA library representation. PCR reactions were then pooled for each sample and column pfrified with QIAGEN PCR purification kit. PCR products were analyzed on an agarose gel, and the DNA band of expected size was excised and purified. Miseq 250-bp paired-end sequencing (Illumina) was performed. For data analysis, FastQ files obtained after MiSeq sequencing were demultiplexed using the MiSeq Reporter software (Illumina). Paired reads were trimmed and filtered using the CLC Genomics Workbench (Qiagen) and matched against sgRNA sequences within the library. Read counts for each sgRNA were normalized against total read counts across all samples. Mageck method were used for differential analysis for sgRNA and Gene ranking (Li et al., 2014c; Wang et al., 2019). A *P* < 0.05 was considered to be statistically significant.

### HUDEP-2 cell culture and induced maturation

Mycoplasma-free HUDEP-2 cells were cultured as described (Kurita et al., 2013). Immature cells were expanded in the StemSpan serum-free medium (SFEM; Stem Cell Technologies) supplemented with 1 µM dexamethasone, 1 µg/mL doxycycline, 50 ng/mL human stem cell factor (SCF), 3 units/mL erythropoietin (EPO), and 1% penicillin–streptomycin. To induce erythroid maturation, HUDEP-2 cells were cultured in a differentiation medium composed of IMDM base medium (Invitrogen) supplemented with 2% FBS, 3% human serum albumin, 3 units/mL EPO, 10 µg/mL insulin, 1000 µg/mL holo-transferrin, and 3 units/mL heparin. Erythroid differentiation and maturation were monitored by flow cytometry, using FITC-conjugated anti-CD235a (BD Biosciences, clone GA-R2, #561017), APC-conjugated anti-Band3 (gift from Xiuli An Lab in New York Blood Center), and VioBlue-conjugated anti-CD49d (Miltenyi, clone MZ18-24A9, #130-099-680).

### CRISPR/Cas9–mediated genome editing of HUDEP-2 cells

The sgRNA sequences were selected from the CRISPR library, generated as oligonucleotides. After annealing, construct was cloned into the *Bbs*I or *Bsm*BI site of the pXPR_003 vector. Lentivirus supernatant were prepared from 293T cells. For cell pool genome editing, HUDEP-2 cells stably expressing Cas9 were transduced with lentiviral vector (pXPR_003) encoding individual sgRNAs. Cells were incubated for 7–10 days with 10 µg/mL blasticidin and 1 µg/mL puromycin to select for transduction with sgRNA and Cas9 vectors, respectively. On-target insertion/deletion mutations were characterized by PCR, followed by next-generation sequencing or TIDE-seq analysis from Sanger sequencing datasets.

### Cell lysates and immunoblot analysis

Cells were suspended in Thermo Scientific Pierce IP Lysis Buffer (ThermoFisher #87787) supplemented with 1 mM phenylmethylsulfonyl fluoride, and 1:500 protease inhibitor cocktail (Sigma–Aldrich). Proteins were resolved on polyacrylamide gels (BioRad), transferred to a PVDF membrane, and incubated in blocking buffer (5% milk in TBST). Antibody staining was visualized using the Odyssey CLx Imaging System.

### (Phospho)proteome sample preparation for MS analysis

All MS experiments were performed in biological quadruplicates. Cell pellets were lysed in SDC buffer (4% Sodium deoxycholate in 100 mM Tris pH 8.5) and heated for 5 min at 95°C. Lysates were cooled on ice and sonicated. Protein concentration was determined by Tryptophan assay as described previously (Kulak et al., 2014). We later made samples up to the equal amounts (120 μg) and in the same volumes (270 μl) and reduced disulphide bonds and carbamidomethylate cysteine residues by adding TCEP and 2-Chloroacetamide to the final volumes of 10 mM and 40 mM, respectively, for 5 min at 45°C. Next, samples were transferred into a 96-deep well plate (DWP). Protein was subsequently digested by the addition of 1:100 LysC and Trypsin overnight at 37°C with agitation (1,500 rpm). Next day, 20 μg of protein material was aliquoted and processed using an in-StageTip (iST) protocol as described previously (Kulak et al., 2014). After peptide clean-up step, concentration was estimated by UV spectrometry and approximately 500 ng was used for single shot DIA analysis. Furthermore, ∼10ug of clean peptides were fractionated using the high-pH reversed-phase ‘Spider fractionator’ into 8 fractions as described previously to generate deep proteomes to build spectral library (Kulak et al., 2017).

The rest of digested peptides (∼100 μg) in the DWP were used for phophospeptide enrichment using the EasyPhos workflow as described previously (Humphrey et al., 2015; Humphrey et al., 2018). After mixing peptides with Isopropanol and EP enrichment buffer (48% TFA, 8 mM KH2PO4), they were enriched with 5mg of TiO2 beads which were prepared at a concentration of 1 mg/μl in loading buffer (6% TFA/80% ACN (vol/vol)) and incubated at 40°C with shaking (2,000 rpm) for 5 min. Afterwards, the phosphopeptide containing TiO2 beads were further washed with 4ml wash buffer (5% TFA/60% ISO (vol/vol)), and treated with elution buffer (40% ACN, 15% NH4OH). Eluted phosphopeptides were concentrated in a SpeedVac for 20min at 45 °C, during which process the volatile salts such as NH4OH and ABC are removed. The samples were then desalted using StageTips loaded with SDB-RPS discs, and again concentrated in a SpeedVac until dry. 6 μl MS loading buffer (0.2% TFA/2% ACN (vol/vol).) was added to the samples, which were then sonicated for 5 min in a bath sonicator.

### Liquid chromatography-MS analysis

Nanoflow LC-MS/MS measurements were carried out on an EASY-nLC 1200 system (ThermoFisher Scientific) combined with the latest generation linear quadrupole Orbitrap instrument (Q Exactive HF-X) coupled to a nano-electrospray ion source (Thermo Fisher Scientific). We always used a 50 cm HPLC column (75 μm inner diameter, in-house packed into the tip with ReproSil-Pur C18-AQ 1.9 μm resin (Dr. Maisch GmbH)). Column temperature was kept at 60°C by a Peltier element containing in-house developed oven.

500 ng peptides were analyzed with a 100 min gradient. Peptides were loaded in buffer A (0.1% formic acid (FA) (v/v)) and eluted with a linear 80 min gradient of 5-30% of buffer B (80% acetonitrile (ACN) plus 0.1% FA (v/v)), followed by a 4 min increase to 60% of buffer B and a 4 min increase to 95% of buffer B, and a 4 min wash of 95% buffer B at a flow rate of 300 nl/min. Buffer B concentration was decreased to 4% in 4 min and stayed at 4% for 4 min.

For the analysis of the fractions to build the project-specific spectral library, the instrument was operated in the DDA mode (Top12). The resolution of the Orbitrap analyzer was set to 60,000 and 15,000 for MS1 and MS2, with a maximum injection time of 20 ms and 60 ms, respectively. The mass range monitored in MS1 was set to 300–1,650 m/z. The automatic gain control (AGC) target was set to 3e6 and 1e5 in MS1 and MS2, respectively. The fragmentation was accomplished by higher energy collision dissociation at a normalized collision energy setting of 27%. Dynamic exclusion was 20 sec.

For single shot samples, the instrument was operated in the DIA mode. Every MS1 scan (350 to 1650 m/z, 120,000 resolution at m/z 200, AGC target of 3e6 and 60 ms injection time) was followed by 33 MS2 windows ranged from 300.5 m/z (lower boundary of first window) to 1649.5 m/z (upper boundary of 33rd window). This resulted in a cycle time of 3.4 s. MS2 settings were an ion target value of 3 × 106 charges for the precursor window with an Xcalibur-automated maximum injection time and a resolution of 30,000 at m/z 200. The fragmentation was accomplished by higher energy collision dissociation with stepped collision energies of 25.5, 27 and 30%. The spectra were recorded in profile mode. The default charge state for the MS2 was set to 3. Data were acquired with Xcalibur 4.0.27.10 and Tune Plus version 2.1 (Thermo Fisher).

Phosphopeptides were analyzed with a 100 min gradient. Peptides were loaded in buffer A (0.1% formic acid (FA) (v/v)) and eluted with a linear 60 min gradient of 3-19 of buffer B (80% acetonitrile (ACN) plus 0.1% FA (v/v)), followed by a 30 min increase to 41% of buffer B and a 5 min increase to 90% of buffer B, and a 5 min wash of 90% buffer B at a flow rate of 350 nl/min. The instrument was operated in the DDA mode (Top10). The resolution of the Orbitrap analyzer was set to 60,000 and 15,000 for MS1 and MS2, with a maximum injection time of 120 ms and 50 ms, respectively. The mass range monitored in MS1 was set to 300–1,600 m/z. The automatic gain control (AGC) target was set to 3e6 and 1e5 in MS1 and MS2, respectively. The fragmentation was accomplished by higher energy collision dissociation at a normalized collision energy setting of 27%. Dynamic exclusion was 30 sec.

### MS data analysis

The fractions (DDA) and the single shot samples (DIA) were used to generate a DDA-library and direct-DIA-library, respectively, which were combined into a hybrid library in Spectromine version 1.0.21621.8.15296 (Biognosys AG). The hybrid spectral library was subsequently used to search the MS data of the single shot samples in Spectronaut version 12.0.20491.9.26669 (Biognosys AG) for final protein identification and quantification. All searches were performed against the Human UniProt FASTA database (2017, X entries). Carbamidomethylation was set as fixed modification and acetylation of the protein N-terminus and oxidation of methionine as variable modifications. Trypsin/P proteolytic cleavage rule was used with a maximum of two miscleavages permitted and a peptide length of 7-52 amino acids. When generating the spectral library generation, minimum and maximum of number of fragments per peptide were set to 3 and 6, respectively. A protein and precursor FDR of 1% were used for filtering and subsequent reporting in samples (q-value mode).

For the phosphoproteome, raw MS data were processed using MaxQuant version 1.6.2.10 (Cox and Mann, 2008; Cox et al., 2011) with an FDR < 0.01 at the peptide and protein level against the Human UniProt FASTA database (2017). Enzyme specificity was set to trypsin, and the search included cysteine carbamidomethylation as a fixed modification and N-acetylation of protein and oxidation of methionine and phosphorylation (SYT) as variable modifications. Up to two missed cleavages were allowed for protease digestion, and peptides had to be fully tryptic.

### Bioinformatics data analysis

We mainly performed data analysis in the Perseus (version 1.6.0.9) (Tyanova et al., 2016), Microsoft Excel and data visualized using GraphPad Prism (GraphPad Software) or RStudio (https://www.rstudio.com/). Apart from coefficient of variation, log2-transformed protein intensities were used for further analysis. Coefficients of variations were calculated for raw protein intensities between replicates individually. Phosphopeptides that were identified in the decoy reverse database were not considered for data analysis. Both data sets were filtered to make sure that identified proteins and phosphopeptides showed expression in all biological triplicates of at least one differentiation stage and the missing values were subsequently replaced by random numbers that were drawn from a normal distribution (width=0.3 and down shift=1.8). PCA analysis of differentiation stages and biological replicates was performed as previously described in (Deeb et al., 2015). Multi-sample test (ANOVA) for determining if any of the means of differentiation stages were significantly different from each other was applied to both mRNA and protein data sets. For truncation, we used permutation-based FDR which was set to 0.05 in conjunction with an S0-parameter of 0.1. For hierarchical clustering of significant proteins, median protein abundances of biological replicates were z-scored and clustered using Euclidean as a distance measure for row clustering. Gene ontology (GO) enrichments in the clusters were calculated by Fischer’s exact test using Benjamini-Hochberg false discovery rate for truncation, setting a value of 0.02 as threshold. Mean log2 ratios of biological triplicates and the corresponding p-values were visualized with volcano plots. We chose a significance cut-off based on a FDR<0.05 in volcano plots.

### Copy number calculation

Intensities were converted to copy number estimations using the proteomic ruler (Wisniewski et al., 2014). The proteomic ruler plug-in v.0.1.6 was downloaded from the Perseus plugin store, for use with Perseus version 1.5.5.0. Protein intensities were filtered for 100% data completeness in at least one stage. Protein groups were annotated with amino acid sequence and tryptic peptide information for the leading protein ID, using the .FASTA file used for processing data. Copy numbers were estimated using the following settings; averaging mode – ‘All columns separately’, molecular masses - ‘average molecular mass’, scaling mode – “Histone proteomic ruler’, ploidy ‘2’, total cellular protein concentration – ‘200 g/l’.

### Selection of marker proteins for cell type deconvolution

To compare the ability of different marker proteins to separate an in silico generated mixture population of cells in different differentiation stages we started from the quantitative protein matrix. Six different sets of marker proteins were selected: ‘sorting’ markers, ‘known’ markers, ‘cluster’ markers, ‘SLC’ markers, ‘combined’ markers and a set of randomly selected 20 proteins (‘any20’). All marker sets (except of ‘any20’) were filtered for an ANOVA q-value < 0.01. For the cluster markers, the top three most significant (smallest ANOVA q-value) proteins for each cluster were selected. The combined markers contain all proteins from the sorting markers, known markers, cluster markers and SLC markers. The ‘any20’ markers were selected by randomly picking 20 proteins from the unfiltered protein list, excluding proteins included in any of the other marker lists.

### Generation of a cell type specific signature matrix

A signature matrix was generated for each of the six marker sets. Only two out of the four cell type replicates, replicate 2 and 4, were used for generating the signature matrices. The signature matrices contain averaged, non-logged intensity values of each marker protein.

### Generation of in silico mixture populations

The two replicates that were not used for the generation of the signature matrix (replicates 1 and 3) were subsequently used for creating in silico mixture populations. We generated mixtures for the intensity averages of replicates 1 and 3, but also for each replicate separately.

The ratios for mixing the different cell types were determined by randomly picking 500 combinations of 5 values (corresponding to the 5 cell types) that add up to 1:

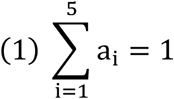

The mixture intensity I_mix_of each protein k was then determined by summing up the intensity of protein k in cell type 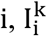, multiplied by the fraction of cells from cell type a_i_:

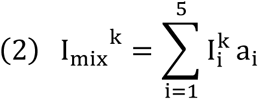

### Deconvolution of the mixture populations

Using the mixture intensities I_mix_ of each marker protein k (Eq. 2) as well as the signature matrix, we set out to estimate the fractions of cells contributed by each of the five cell types 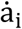. The closer the estimations 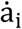 are to the true mixing ratios a_i_, the better the marker set is for deconvoluting and differentiating the five cell types.

Writing equation 2 in matrix form when several proteins are evaluated at the same time, the model expands to the following:

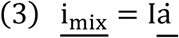

Here, <u>i_mix_</u> is a vector of mixture intensities for each evaluated marker protein k, I is the signature matrix containing the intensities of each evaluated marker protein k (rows) in each cell type i (columns), and 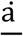 is the vector with the fractions of cells in each cell type i that we aim to estimate.

To estimate 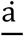, we can solve the linear equation system (Eq. 3) by using a minimum least squares optimization (python scipy.optimize.minimize). Boundary conditions set the minimum possible values of 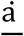 to zero in order to avoid negative values. After an initial estimation of 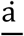, only the top 90^th^ percentile of marker proteins for which the estimates fit best are kept for solving the linear equation system (Eq. 3) in a second iteration. The estimated vector of fractions, 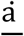, is finally normalized by the Manhattan (L1) norm.

### Evaluation of the cell type deconvolution

To evaluate the results of the deconvolution step, we implemented a weighted error metric to estimate how close the estimated ratios 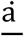 are to the true ratios <u>a</u>. Here, mistakes in assigning neighboring cell types (e.g. between P3 and P4) contribute less to the overall error than mistakes between far distant cell types (e.g. Progenitors and P5). In addition to evaluating the six sets of protein markers, three controls were generated: ‘random’ uses a random ratio estimation; ‘uniform’ assumes a uniform ratio estimation 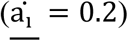; and ‘center’ assumes that all cells are of type P3 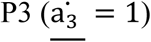.

## SUPLEMENTARY TABLES

**Table S1**. DIA proteome_all quantified proteins with copy numbers.xlsx

**Table S2**. Dynamically expressed SLCs and characterized stage-specific markers

**Table S3**. DDA phosphoproteome_all quantified phosphosites .xlsx

**Table S4**. Expansion hits.xlsx

**Table S5**. Maturation hits_Band3 high vs low.xlsx

## SUPPLEMENTARY FIGURES AND LEGENDS

**Figure S1, Related to Figure 1.**
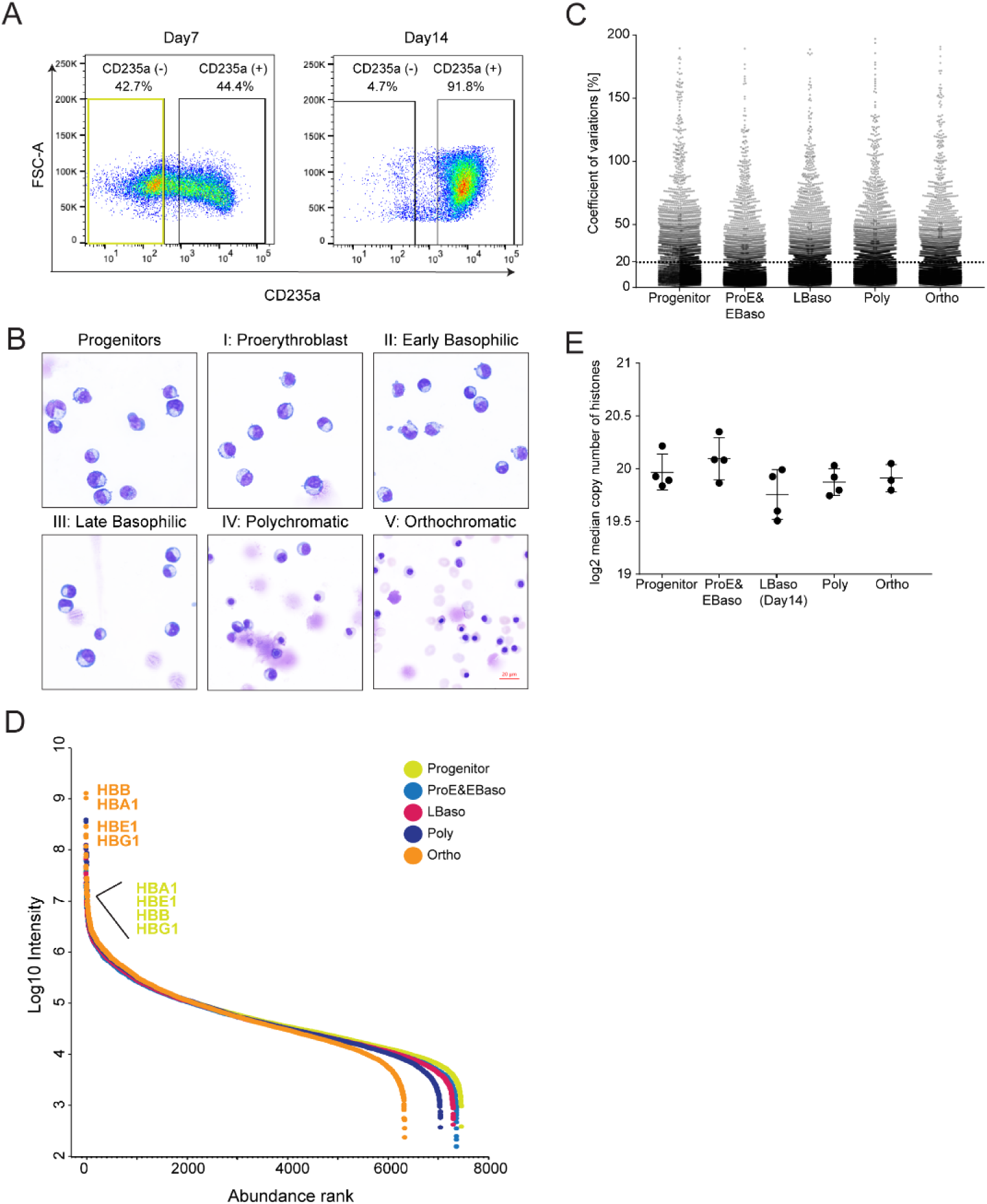
(A) FACS gating/sorting regime to enrich for CD235a^-^ progenitor population. (B) Characterization of the differentiation stages in culture. May-Grünwald-Giemsa staining of erythroid cells is shown. Scale bar, 20 μM. (C) Coefficient variations (CVs) of four biological replicates for each protein were calculated in all stages to show the reproducibility of our system. Dashed line shows the cutoff line of 20% CV. (D) Cumulative protein abundance and dynamic range in five differentiation stages. Hemoglobin subunits (HBB, HBA1, HBE1 and HBG1) are labeled as progenitor (yellow) and Ortho (orange) stages. (E) Estimated median copy numbers of histones per cell across all measured stages.

**Figure S2, Related to Figure 2.**
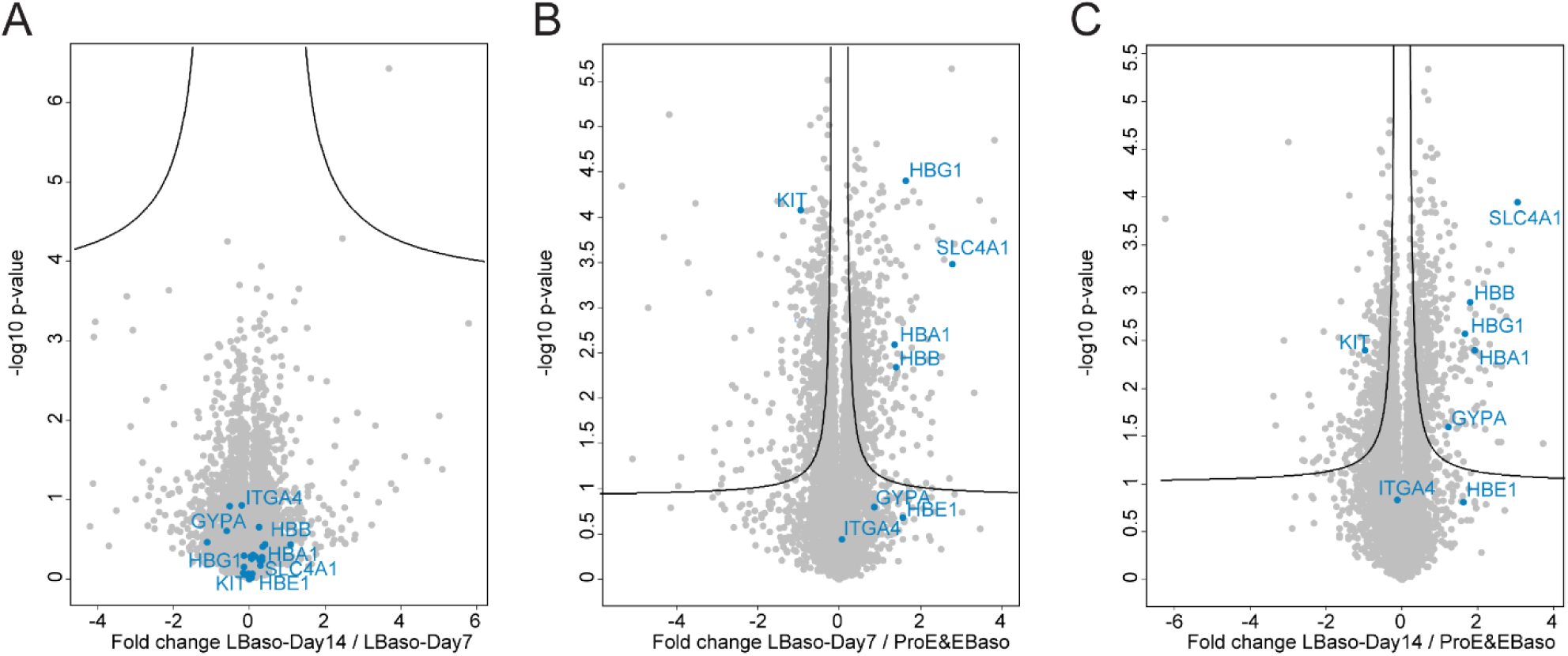
(A-C) Volcano plots of the (-log10) p-values vs. the log2 protein abundance differences between LBaso-Day7 and LBaso-Day14 (left), LBaso-Day7 and ProE-EBaso (middle), and LBaso-Day14 and ProE-EBaso (right) with the significance lines (FDR < 0.05 and S0=0.1). Selected marker proteins are labeled in blue.

**Figure S3, Related to Figure 2.**
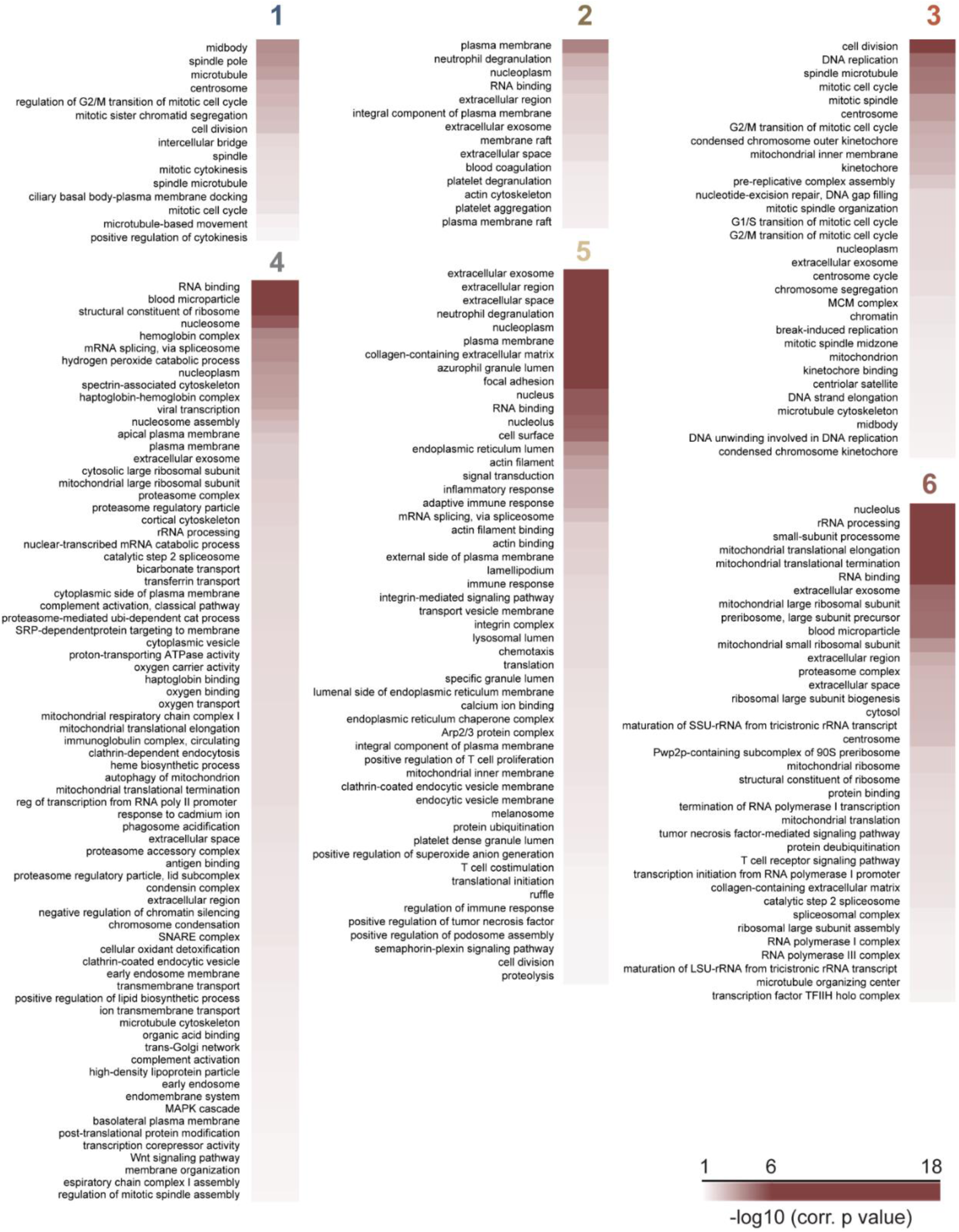
Gene Ontology (GO) enrichment analysis of six clusters of significant proteome shown in Figure 2A was performed using Fischer’s exact test. 2% threshold was applied to Benjamini-Hochberg FDR to determine the significance.

**Figure S4, Related to Figure 2.**
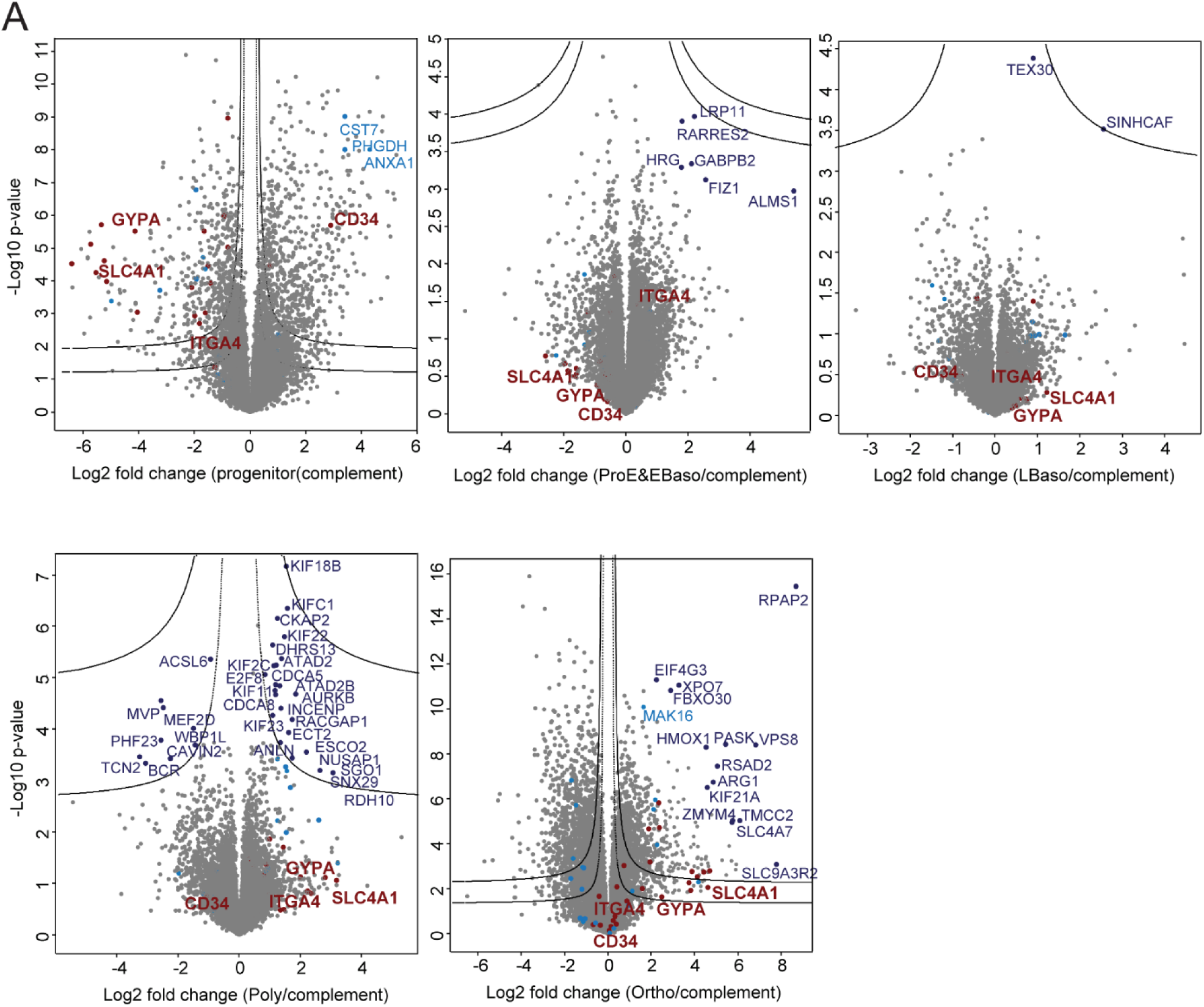
Hawaii plots that overlay all volcano plots of protein enrichments in a specific stage over all other stages plotted against corresponding p-values. Two cut-off lines were placed graphically, defining two confidence classes with FDRs of 0.01 and 0.05 (S0=0.1). Sorting and cluster markers, and selected outliers are labeled in dark red, light blue and dark blue, respectively.

**Figure S5, Related to Figure 5.**
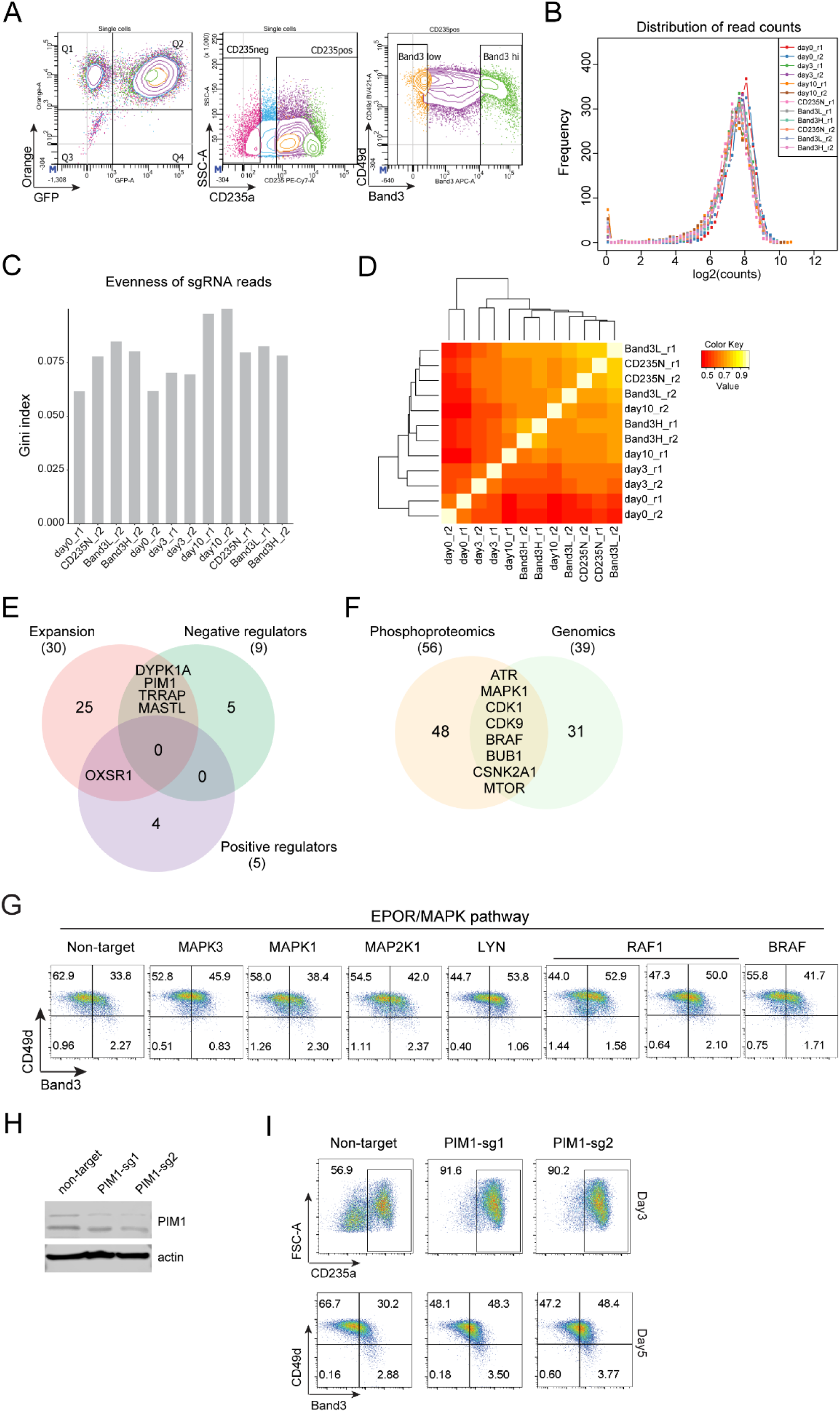
(A) Flow cytometry strategy based on GFP, CD235a, CD49d and Band3 to determine the significant hits from the genome-scale CRISPR-Cas9 screen. (B) Histogram of the sgRNA distribution in each sample in the CRISPR-Cas9 kinase screen. (C) Evenness of the sgRNA reads in each sample in the CRISPR-Cas9 kinase screen. (D) Correlation based reproducibility analysis between replicates in the CRISPR-Cas9 kinase screen. High and low correlation values are denoted in yellow and orange, respectively. (E) Overlap between expansion hits and positive or negative maturation regulators. (F) Overlap of kinases whose activities were inferred by stage-specific substrate profiling from phosphoproteomics in Figure 4F and the genomic CRISPR-Cas9 kinase screen. (G) FACS analysis of Band3 vs CD49d expressions after three days of differentiation in negative control non-targeting and individual sgRNA targeting MAPK1, MAPK3, MAP2K1, LYN, RAF1, BRAF cells. (H) Immunoblot analysis with indicated antibodies of whole cell lysates of non-targeting and PIM1-targeting HUDEP-2 cells. Actin served as protein loading control. (I) Non-targeting and PIM1-targeting (PIM1-sg1 and PIM1-sg2) HUDEP-2 cells were subjected to maturation protocol, followed by flow cytometry of CD235a and Band3/CD49d expressions after three and five days, respectively.

## Notes

### Competing Interest Statement

The authors have declared no competing interest.

